# A transient apical extracellular matrix relays cytoskeletal patterns to shape permanent acellular ridges on the surface of adult *C. elegans*

**DOI:** 10.1101/2021.12.28.474392

**Authors:** Sophie S. Katz, Trevor J. Barker, Hannah M. Maul-Newby, Alessandro P. Sparacio, Ken C.Q. Nguyen, Chloe L. Maybrun, Alexandra Belfi, Jennifer D. Cohen, David H. Hall, Meera V. Sundaram, Alison R. Frand

**Affiliations:** Department of Biological Chemistry, David Geffen School of Medicine, University of California, Los Angeles; Department of Genetics, University of Pennsylvania Perelman School of Medicine; Department of Neuroscience, Albert Einstein College of Medicine

**Keywords:** apical constriction, epidermal morphogenesis, cuticle, alae, molting, Zona-Pellucida domain proteins

## Abstract

Apical extracellular matrices can form protruding structures such as denticles, ridges, scales, or teeth on the surfaces of epithelia. The mechanisms that shape these structures remain poorly understood. Here, we show how the actin cytoskeleton and a provisional matrix work together to sculpt acellular longitudinal alae ridges in the cuticle of adult *C. elegans*. Transient actomyosin-dependent constriction of the underlying lateral epidermis accompanies deposition of the provisional matrix at the earliest stages of alae formation. Actin is required to pattern the provisional matrix into longitudinal bands that are initially offset from the pattern of longitudinal actin filaments. These bands appear ultrastructurally as alternating regions of adhesion and separation within laminated provisional matrix layers. The provisional matrix is required to establish these demarcated zones of adhesion and separation, which ultimately give rise to alae ridges and their intervening valleys, respectively. Provisional matrix proteins shape the alae ridges and valleys but are not present within the final structure. We propose a morphogenetic mechanism wherein cortical actin patterns are relayed mechanically to the laminated provisional matrix to set up distinct zones of matrix layer separation and accretion that shape a permanent and acellular matrix structure.

## INTRODUCTION

Apical extracellular matrices (aECMs) line all epithelial surfaces in contact with the environment. These aECMs vary in composition, but typically contain a mix of proteins, carbohydrates and lipids that are organized into recognizable layers. Some aECMs are soft and gel-like, but others form more rigid structures with characteristic shapes, such as the hook-like denticles or cuticle ridges of insects and nematodes (Cox, 1981; Fernandes et al., 2010) or the scales on butterfly wings (Lloyd and Nadeau, 2021). Examples of aECM in humans include mucin- and proteoglycan-rich linings within many tube lumens (Gaudette et al., 2020; Johansson et al., 2013; Whitsett et al., 2015); the tectorial membrane, a flexible aECM sheet that relays sound waves within the inner ear (Sellon et al., 2019); hair, an amalgamation of keratinized cells and extracellular macromolecules (Harland and Plowman, 2018); and tooth enamel, a composite of calcium phosphate minerals and matrix proteins (Moradian-Oldak and George, 2021). Mutations that affect component matrix proteins cause various disease phenotypes (Gaudette et al., 2020; Schaeffer et al., 2021; Sellon et al., 2019; Whitsett et al., 2015). Despite the widespread functional and medical significance of the aECM, the cellular and molecular mechanisms that sculpt apical matrices are not well understood.

In some cases, the shape of an aECM structure is molded at least in part by the shape of the underlying epithelium at the time of matrix deposition. For example, denticles and taenidial ridges on insect cuticles originate as actin-based cellular protrusions that subsequently become coated with aECM (Fernandes et al., 2010; Hannezo et al., 2015; Ozturk-Colak et al., 2015). The cellular protrusions eventually withdraw, leaving the rigid aECM structures in place. Differences in matrix composition affect not only denticle or taenidia shape but also the apical domain architecture within the underlying cells, suggesting mechanical connections among the aECM, apical junctions, and the actin cytoskeleton. However, these mechanical links between aECM and the cytoskeleton have yet to be fully elucidated. Furthermore, some complex aECM structures such as nematode alae are not obviously associated with cellular protrusions (Cox, 1981; Sapio et al., 2005), raising the question of how such acellular structures are shaped.

The *C. elegans* body cuticle is a multi-layered aECM composed mainly of collagens (Page and Johnstone, 2007). *C. elegans* sheds and replaces its cuticle by molting to progress between each of its four larval stages and to enter adulthood (Knight et al., 2002; Singh and Sulston, 1978; Sulston and Horvitz, 1977). The cuticle of each stage is unique in structure: Longitudinal acellular ridges or “alae” form above the lateral (seam) epidermis in L1s, dauer larvae, and adults, but not in the intervening L2, L3, or L4 stages (Fig. 1) (Cox, 1981). Therefore, alae patterns must be generated *de novo*, rather than propagated from one life stage to the next across the molts. The number, size, and shape of alae ridges also varies among L1s, dauer larvae, and adults (Cox, 1981). The basis of these stage-specific morphologies is unclear, except for the differential requirements for specific aECM proteins, including Zona Pellucida (ZP)-domain cuticulin proteins and various provisional matrix components (Cohen et al., 2019; Flatt et al., 2019; Forman-Rubinsky et al., 2017; Muriel et al., 2003; Sapio et al., 2005; Sebastiano et al., 1991).

**Figure 1.**
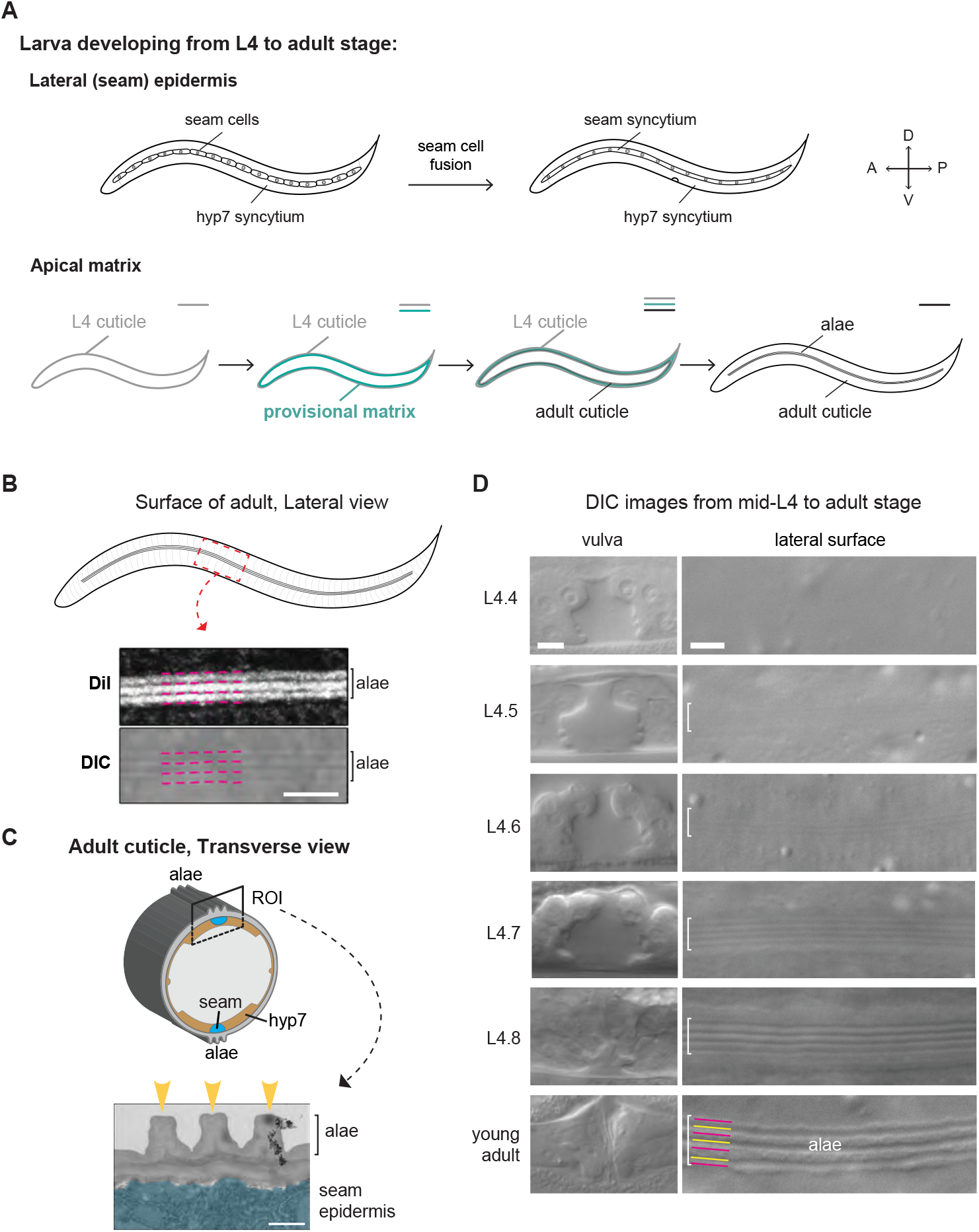
Timeline of adult alae formation. A) L4-to-Adult development. Top schematics show anatomy of the seam and hyp7 syncytia. Bottom schematics show apical matrices. The provisional matrix (teal) is secreted beneath the L4 cuticle (gray) prior to, or during, synthesis of the new adult cuticle (black). B) Lateral view of the adult cuticle. Micrographs show adult alae as visualized by DiI staining and DIC. Magenta lines indicate dark bands corresponding to valleys in both the DiI and DIC images. Scale bar: 5 μm. C) Transverse view of adult. Cartoon (top) shows how alae are positioned relative to the seam and hyp7 syncytia. Transmission electron micrograph (bottom) shows the three alae ridges (yellow arrowheads). The underlying seam syncytium is false colored in blue. Scale bar: 500 nm. D) When viewed by DIC imaging, alae on the developing adult cuticle gradually became visible underneath the L4 cuticle. Brackets indicate regions above seam syncytia where longitudinal ridges could be detected. N2 animals were grown at 20°C and staged based on vulva tube morphology (Mok et al., 2015). Images are representative of at least n=10 animals per stage. Scale bars: 5 μm.

Actomyosin-dependent forces and cuticle matrix buckling have been indirectly implicated in alae formation at the L1 and dauer stages, but the specific steps involved have not been characterized in detail. L1 cuticle formation occurs in the context of actomyosin-driven seam cell apical constriction, which elongates the embryo and narrows the entire body shape (Priess and Hirsh, 1986; Zhang et al., 2011). Similarly, dauer larvae are radially-constricted compared to the preceding larval stage (Flatt et al., 2019; Sebastiano et al., 1991; Singh and Sulston, 1978). Cuticulin mutants that lack alae at one or both of these stages also have wider seam cells and a dumpy body shape (Sapio et al., 2005). Based on these observations, Sapio et al. (2005) proposed that stage-specific ZP proteins somehow generate or distribute constrictive forces that fold the developing cuticle into alae ridges over the narrowing seam cells. However, this “cuticle bending” model was not further tested experimentally and also did not explain how alae form in the adult, where overall volume increases while the general body shape remains similar to that of the preceding L4 stage (Fig. 1A).

Recent work has shown that a ZP-rich provisional matrix precedes formation of each *C. elegans* cuticle (Cohen and Sundaram, 2020). In the embryo, the first cuticle is synthesized beneath a provisional matrix termed the embryonic sheath (Costa et al., 1997; Priess and Hirsh, 1986; Vuong-Brender et al., 2017). During the molt cycle, epidermal cells secrete a new provisional apical matrix beneath each pre-molt cuticle before synthesis of the post-molt cuticle (Fig. 1A) (Forman-Rubinsky et al., 2017; Gill et al., 2016). Components of the provisional matrix are removed before or along with the pre-molt cuticle. This transient provisional matrix may help maintain tissue and body shape during the molting process. The provisional matrix also influences the organization of the subsequent cuticle and shapes the alae of the L1, dauer and adult stages, potentially by acting as a scaffold for further matrix deposition (Forman-Rubinsky et al., 2017; Gill et al., 2016).

Here, we examined how the actin cytoskeleton and the provisional matrix work together to sculpt the alae of adult *C. elegans*. Transient actomyosin-dependent narrowing of the seam surface accompanies deposition of the provisional matrix at early stages of alae formation, but it does not appear to buckle the apical membrane. Instead, longitudinal actin filament bundles (AFBs) at the seam cortex align with ultrastructural delaminations and future valleys that flank alae ridges. Actin is required to pattern the provisional matrix into longitudinal bands, and the provisional matrix is required to establish demarcated zones of matrix layer adhesion and separation which ultimately give rise to alae ridges and their intervening valleys, respectively. We propose a morphogenetic mechanism wherein cortical actin patterns are relayed mechanically to the laminated provisional matrix to set up distinct zones of matrix layer separation and accretion, thereby shaping a permanent and acellular matrix structure.

## RESULTS

### Morphogenesis of the adult-stage alae begins midway through the L4 stage

The *C. elegans* epidermis consists of cells and multinucleate syncytia that together synthesize most of the external cuticle (Podbilewicz and White, 1994; Sulston and Horvitz, 1977). The lateral seam and adjacent hyp7 syncytium are the two largest tissues and are connected by apical-lateral junctions analogous to those found in vertebrates and insects (Pasti and Labouesse, 2014) (Fig. 1A). The seam cells undergo stem cell-like asymmetric divisions early in each larval stage: anterior daughters fuse with hyp7, while posterior daughters remain in place and reconnect. During L4, seam cells exit the cell-cycle, fuse into bilateral syncytia, and ultimately synthesize alae - three key events that mark the L4-to-adult transition (Sulston and Horvitz, 1977).

Adult alae consist of three longitudinal ridges that decorate the cuticle overlying the seam (Cox, 1981) (Fig. 1). Alae ridges stain prominently with the lipophilic fluorescent dye DiI (Schultz and Gumienny, 2012) (Fig. 1B). Alae also can be seen with Differential Interference Contrast (DIC) microscopy as alternating dark and light stripes, which we confirmed correspond to the three ridges and four flanking valleys, respectively (Fig. 1B). As viewed by transmission electron microscopy (TEM), alae are acellular structures and protrude approximately 0.5 microns above the rest of the cuticle surface (Fig. 1C).

We were able to visualize the timing of adult alae formation using DIC microscopy of L4 animals staged based on developing vulva tube morphology (Fig. 1D). Longitudinal stripes on the lateral surface first became visible at stages L4.5-L4.6. The stripes were initially subtle but gradually became more prominent at later stages. Thus, morphogenesis of the alae began well before the L4-to-adult molt and appeared to be a gradual rather than abrupt process.

### Actomyosin filaments in both the seam and hyp7 shape the adult-stage alae

To test the role of the actin cytoskeleton in forming alae, we used bacterial-mediated RNA-interference (RNAi) to silence actin genes in developing larvae and later examined the lateral surfaces of young adults by DIC microscopy and staining with DiI.

Five *C. elegans* genes encode actin monomers. During the L4 stage, epidermal cells and syncytia express *act-2* most highly, with some evidence for expression of *act-1, −3* and *-4* at other stages (Katsanos et al., 2021; Willis et al., 2006). To simultaneously knock down *act-1, −2, −3* and *-4*, we selected an *act-2*-derived dsRNA trigger complementary to all four transcripts (Methods). Further, we customized and applied an established experimental paradigm to selectively knock down actin in either the seam or hyp7. This system involved tissue-specific expression of RDE-1, the worm homolog of Argonaute, in *rde-1(ne219)* mutants otherwise insensitive to siRNAs (Qadota et al., 2007). This approach bypassed the embryonic and larval lethality associated with systemic *actin(RNAi)* over the full course of development. Waiting until the L2 stage to deliver *actin* dsRNAs also bypassed much of this lethality; attenuated *actin(RNAi)* larvae developed into small adults.

Attenuated *actin(RNAi)* or preferential knockdown of actin in either the seam or hyp7 resulted in patches of fragmented and disorganized adult-stage alae (Fig. 2A,C,D). The most severely affected animals lacked continuous ridges entirely, while others retained the outer dorsal and ventral ridges but showed breaks and misorientations within the central alae ridge, resulting in a “braid-like” appearance. Larger gaps in the alae (> 5 microns long) sometimes flanked the disorganized regions, and in these cases the underlying epidermis was likely disrupted due to earlier defects in seam cell division and reconnection, as previously described (Ding and Woollard, 2017). Because cell division and fusion were not of interest in this study, our approaches were designed to minimize such defects and our analysis prioritized extended regions of alae disorganization over any large gaps in the alae (Fig. 2D, Methods). The abovementioned alae deformities were not observed in *rde-1* null mutants fed *actin* dsRNAs or in *wild-type* animals fed short dsRNAs transcribed from the vector. The fact that knockdowns in either the seam or hyp7 caused similar deformities suggests that actin networks within both syncytia work cooperatively to shape the adult-stage alae.

**Figure 2.**
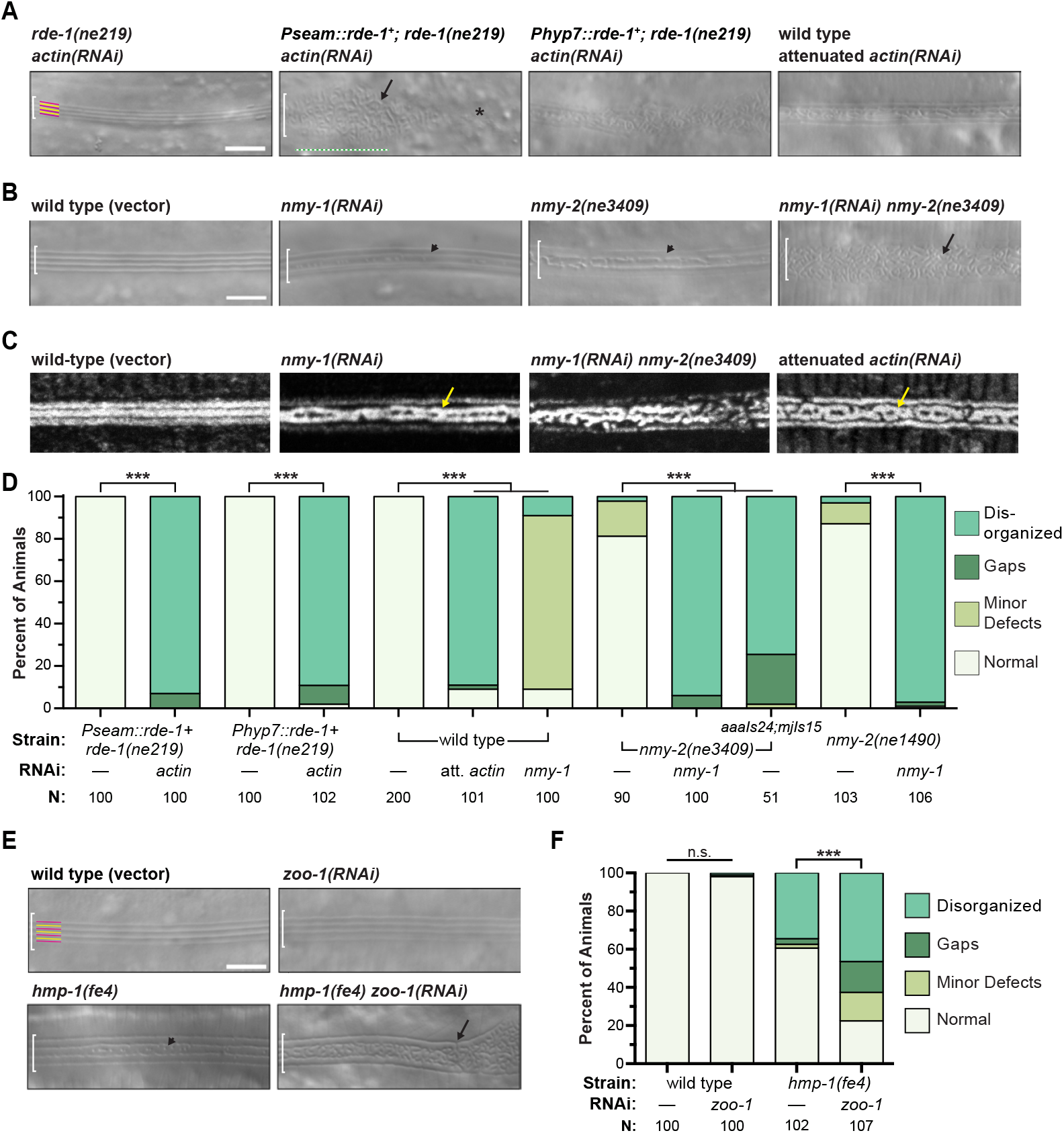
Knockdown of actin, NM II, or AJ components results in alae disorganization. A) Tissue-specific and whole animal actin knockdowns (strains JK537, ARF330, ARF408, N2, 25°C). B) NM II mutants and knockdowns (strains WM179, WM180, 25°C). For both A and B, representative DIC images show structures on the lateral surface of young adults of indicated genotypes. White brackets demarcate the region of interest. Yellow and magenta lines label presumptive ridges and adjacent valleys in normal alae. Arrows point to tortuous structures; arrowheads, minor deformities. Asterisk labels large gap. Green bracket labels severely disorganized region with no obvious ventral boundary. C) DiI staining of the cuticle in actin and NM II knockdowns. D) Prevalence of deformed alae. Values are weighted averages from two independent trials; N: sample size. *** P<0.001 for all pairwise comparisons of prevalence of seemingly normal alae in knockdowns and mock-treated specimens; Fisher’s exact test with Bonferroni’s correction for multiple comparisons. E-F) AJ mutants or knockdowns (strains N2 and PE97, 25°C), and prevalence of deformed alae, as above. ***P < 0.001. Scale bars: 5 μm.

Non-muscle myosin (NM II) often partners with actin to generate morphogenetic forces (Martin and Goldstein, 2014; Munjal and Lecuit, 2014). The *nmy-1* and *nmy-2* genes of *C. elegans* both encode heavy chains of NM II that are 47% identical in primary sequence to human NMHC-IIB and expressed in the epidermis (Piekny et al., 2003). We used *nmy-1(RNAi)* and conditional alleles of *nmy-2* to partially inactivate NM II. Fragmented and disorganized alae were observed on most *nmy-1(RNAi); nmy-2(ts)* double mutants cultivated at restrictive temperature (Fig. 2B-D). In contrast, only minor deformities in the alae were observed in *nmy-1(RNAi)* or *nmy-2(ts)* single mutants, although *nmy-2(ts)* defects were greatly enhanced by expression of an F-actin biosensor and junction marker (see below) (Fig. 2D). The combinatorial effect of *nmy-1(RNAi)* and *nmy-2(ts)* suggests that these paralogs make redundant contributions to a morphogenetic mechanism involving actomyosin-dependent forces.

### Apical Junction (AJ) components that interact with actin networks shape the adult alae

Actomyosin filaments often attach to cell membranes at cell-cell junctions (Martin and Goldstein, 2014; Munjal and Lecuit, 2014). To evaluate the role of AJs in patterning the adult alae, we similarly used RNAi and a hypomorphic allele to knock down key AJ components while larvae developed and then examined the lateral surface of young adults. HMP-1/α-catenin is the actin-binding component of cadherin-catenin complexes (CCCs) that mechanically link the various epidermal cells of *C. elegans* (Costa et al., 1998; Kang et al., 2017; Pasti and Labouesse, 2014). The Zonula Occludens (ZO) homolog ZOO-1 cooperatively recruits actin bundles to AJs (Lockwood et al., 2008). Defective alae were observed on the surface of more than one third of surviving *hmp-1(fe4)* hypomorphic mutants, and these defects were further enhanced by simultaneous knockdown of *zoo-1* (Fig. 2E-F). Thus, genetic manipulations known to impede the transmission of mechanical forces through AJs interfered with morphogenesis of adult-stage alae.

### Transient narrowing of the seam apical cortex precedes alae formation

To investigate the mechanism by which actomyosin networks pattern adult-stage alae, we further characterized the superficial shape of the seam epidermis and organization of cortical actin across the L4 stage. For this purpose, we used AJM-1 fusion proteins to label AJs among the seam and hyp7 syncytia (Fig. 3A,B) (Koppen et al., 2001; Lehrbach et al., 2009). We also constructed and used a highly sensitive sensor for F-actin composing the Calponin homology domain (CH) of human Utrophin (UTRN) tagged with GFP and driven by the seam-specific promoter of *egl-18* (Koh and Rothman, 2001) (Fig. 3A,B). This CH domain binds F-actin selectively and reversibly, and UTRNCH::GFP does not appreciably perturb actin dynamics when expressed at practical levels (Burkel et al., 2007; Moores and Kendrick-Jones, 2000). To achieve fine temporal resolution, we isolated precisely staged transgenic nematodes and imaged them at regular ~1hr intervals (Methods). We also collected images of the vulva to assess animal stage directly (Fig. 1D).

**Figure 3.**
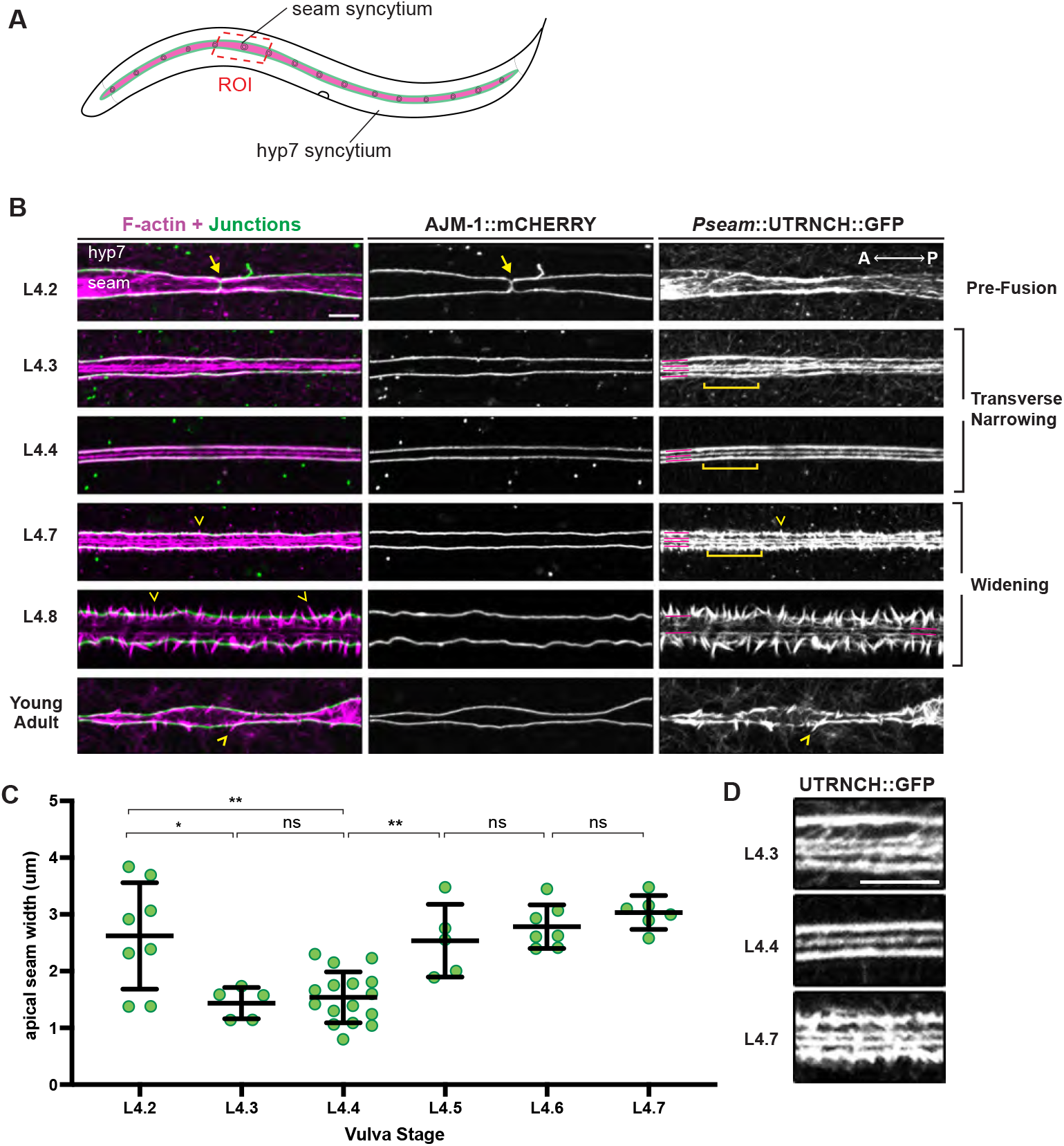
Transient narrowing of seam syncytia and formation of longitudinal AFBs precede the appearance of alae. A) Diagram depicting seam (magenta) and hyp7 syncytia in an L4 larva. Green, apical junctions between syncytia. Red box indicates region of interest (ROI) imaged below. B) Representative confocal projections show UTRNCH::GFP (magenta) and AJM-1::mCHERRY (green) signals captured at the indicated stages (strain ARF404, 25°C). Yellow lines label AFBs along dorsal and ventral seam margins; dashed lines, medial AFBs. Arrows point to AJs between seam (s) cell cousins about to fuse. Chevrons point to spikes of F-actin crossing AJs. C) Quantification of apical width of seam syncytia at indicated stages (strain SU93, 20°C). Values are the distance between AJM-1::GFP marked AJs, representing the mean of 6 measurements per worm. Bars: mean and s.d. *P ≤ 0.05, **P ≤ 0.01; Mann-Whitney test. D) Enlarged ROIs delineated by brackets in (B). Scale bars: 5 μm.

The AJ marker revealed that seam cell fusion occurred immediately following the last round of cell divisions early in the L4 stage, with fusion complete before or during L4.2 (Fig. 3B). Soon after seam fusion, the apical surface of seam syncytia narrowed, appearing most narrow and uniform at L4.3 to L4.4 (Fig. 3B,C). The apical surface of the seam then widened slowly, as the AJs spread apart over the following several hours and animals completed the L4/adult molt (Fig. 3B,C). The observed cell shape changes suggested transient apical constriction on the transverse or dorsal-ventral (D-V) axis prior to initial alae formation, followed by gradual relaxation as the alae form and enlarge.

### Longitudinal AFBs in the seam presage the pattern of adult alae

The seam-specific UTRNCH::GFP marker revealed that striking changes in actin appearance accompanied seam narrowing and relaxation (Fig. 3B,D). At the onset of narrowing (~L4.3), four longitudinal AFBs assembled at the cortex of the seam syncytia; two of these AFBs co-localized with AJM-1::mCHERRY along the dorsal and ventral junctions with hyp7, while the other two AFBs were located more medially. At the narrowest seam stage (L4.3-L4.4), often only three longitudinal AFBs were detected, suggesting the medial AFBs had moved closer together and potentially joined. As the seam widened again (L4.5-L4.7), four longitudinal AFBs were again observed.

Medial AFBs began to disassemble and breaks in the outer junctional AFBs appeared at the L4.7-L4.8 stages (Fig. 3B), after the time that alae first become visible by DIC (Fig. 1D). Spikes of F-actin that apparently crossed the dorsal and ventral junctions also became more prominent at this time. This progressive transition from continuous longitudinal to discontinuous transverse F-actin structures along the margins might reflect a concurrent transition in the net direction of force propagation between the seam and hyp7.

In summary, the transient narrowing of seam syncytia midway through L4 coincided with dynamic reorganization of the bulk of cortical F-actin into three or four longitudinal AFBs. This pattern is not what we would expect for a typical apical constriction process, where AFBs are usually either isotropic or aligned in the direction of tissue narrowing (Koenderink and Paluch, 2018). However, these distinctive arrangements are reminiscent of the adult alae pattern of three longitudinal ridges and four flanking valleys, which first manifest while these AFB patterns are still present.

### Actin assembles into both longitudinal and circumferential AFBs in hyp7

As our RNAi experiments indicated that actomyosin networks in hyp7 syncytia also contribute to morphogenesis of the alae (Fig. 2), we went on to track the distribution of cortical actin in hyp7 across the L4 stage (Fig. 4). For this purpose, we constructed a similar but distinct F-actin sensor comprising UTRNCH tagged with dsRED and driven by the hypodermal-specific promoter of *dpy-7* (Gilleard et al., 1997). This sensor revealed both longitudinal AFBs along the hyp7-seam margins and circumferential filament bundles (CFBs), some of which branched off from the longitudinal bundles (Fig. 4A).

**Figure 4.**
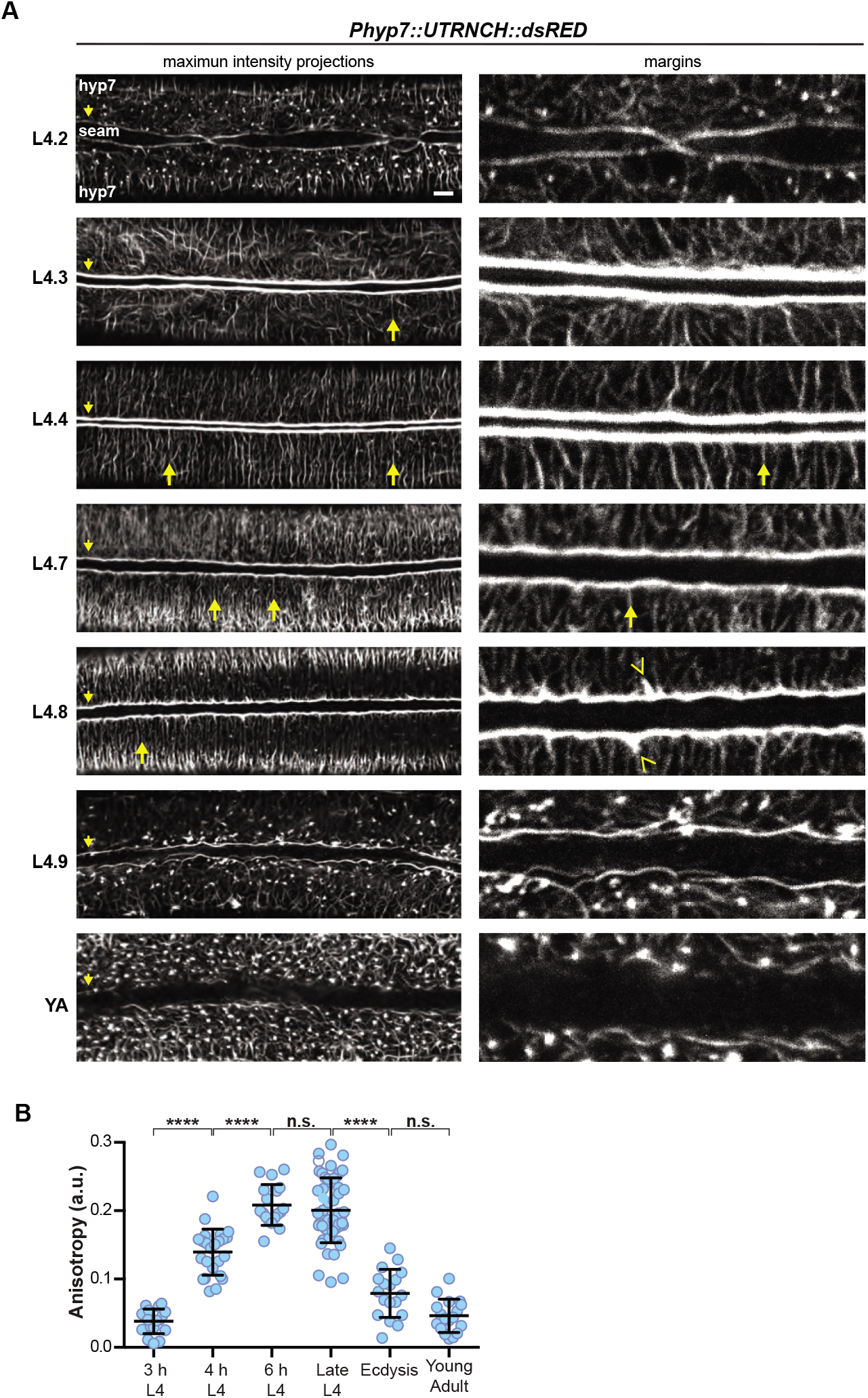
Assembly of both longitudinal and circumferential AFBs in hyp7 precede the appearance of alae. A) Confocal fluorescence maximum-intensity projections (left) show F-actin (UTRNCH::dsRED) detected in the lateral (thick) region of hyp7 syncytia. Digitally enlarged ROIs show F-actin by the lateral margins (strain ARF385, 25°C except for L4.2 specimen, 20°C). Arrowheads point to F-actin at margins. Arrows point to circumferential AFBs. B) Quantitation of anisotropy of F-actin signals in lateral hyp7. Values represent 6 ROIs from 3-10 specimens per timepoint (relative to emergence in L4) or stage. ****P ≤ 0.0001, ^n.s.^P > 0.05; Ordinary one-way ANOVA with Tukey’s correction.

Like seam AFBs, the hyp7 longitudinal AFBs formed immediately following seam cell fusion (Fig 4A,B). These AFBs remained prominent throughout seam narrowing and relaxation (as defined in Fig. 3) and then disappeared at the L4-adult molt. We infer that longitudinal seam and hyp7 AFBs run in parallel along each side of the seam-hyp7 AJs.

The hyp7 CFBs have long been considered a hallmark of molting animals (Costa et al., 1997), but our analysis showed that they assemble earlier than previously thought, and that their appearance changes at the time of seam narrowing (Fig. 4A). Prior to seam narrowing, some whisker-like filaments branched off from the longitudinal bundles at variable angles. Following seam narrowing, CFBs became increasingly anisotropic (aligned in parallel) up until the end of the L4-adult molt, when they collapsed at ecdysis (Fig. 4C).

### Seam narrowing and AFB organization depend on NM II

If AFBs and/or CFBs are contractile structures, we would expect them to associate with NM II (Martin and Goldstein, 2014; Munjal and Lecuit, 2014). To test this, we examined the localization patterns of NMY-1::GFP and NMY-2::GFP expressed from the endogenous loci (Dickinson et al., 2013; Vuong-Brender et al., 2017), and we generated a seam UTRNCH::dsRed reporter and compared the patterns of actin and NMY-2::GFP (Fig. 5A,B). Surprisingly, while UTRNCH::dsRed marked three longitudinal AFBs upon seam narrowing, it differed from UTRNCH::GFP in that, during relaxation, it clearly marked only the two junctional AFBs and only faintly if at all marked the two medial AFBs (Fig. 5A). One difference between GFP and dsRed is that the latter is an obligate tetramer (Baird et al., 2000), so differences in conformation might explain the discrepancy in the patterns seen with these two actin sensors. Whatever the explanation, the differential ability of UTRNCH::dsRed to mark junctional vs. medial AFBs suggests that these AFBs may differ in organization.

**Figure 5.**
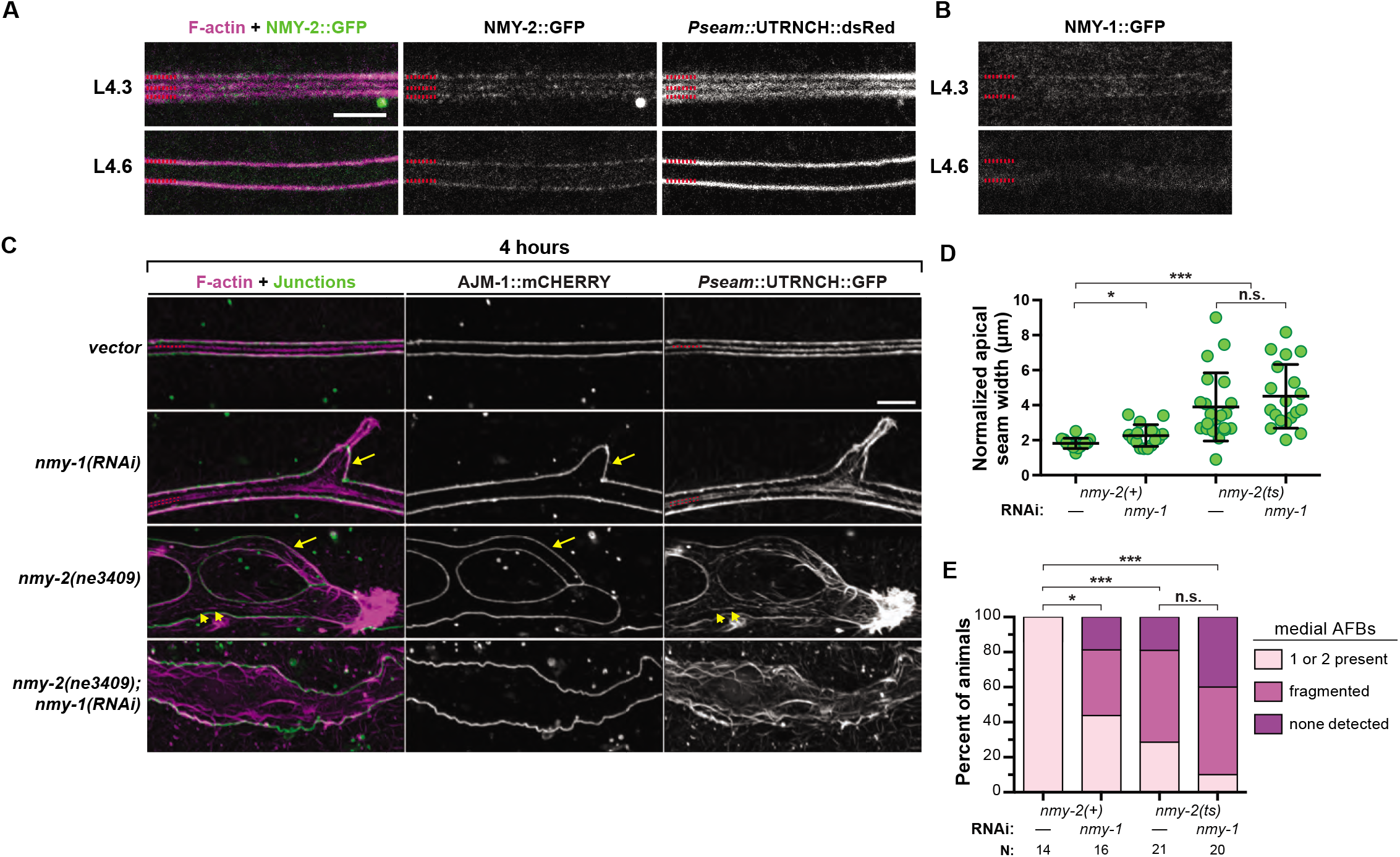
Seam narrowing and coherence of AFBs depend on NM II. **A)** Endogenous NMY-2::GFP aligns with junctional AFBs at the seam-hyp7 margins and with the medial AFB in narrowed (L4.3-L4.4) seam syncytia (strain ARF500 20°C). AFBs were marked by *egl-18::*UTRNCH::dsRed. B) Endogenous NMY-1::GFP also faintly marks the seam-hyp7 margins (Strain ML2540, 20°C). Both A and B show single confocal slices representative of at least n=6 animals per stage, imaged at 20°C. C) NM II knockdown disrupts seam shape and medial AFBs. Representative confocal maximum projections show F-actin and AJs detected near the surface of seam syncytia at L4.4 (strain ARF404, 25°C). Solid lines label junctional AFBs; dashed lines, medial AFBs. Arrows point to protrusions of the seam over hyp7. Asterisks label aggregated F-actin. Arrowheads point to fragmented medial AFBs. Scale bar: 5 μm. D) Quantitation of seam width — values represent area surrounded by AJs normalized to imaged interval. Images and measurements from two independent trials; total sample sizes as indicated. Bars signify mean and s.d. ***P ≤ 0.001, *P ≤ 0.05; unpaired t-tests with Bonferroni’s correction. For C and D, strains expressed *egl-18*::UTRNCH::GFP and AJM-1::mCHERRY.

NMY-1::GFP and NMY-2::GFP puncta aligned along or near the two junctional AFBs at the presumptive seam margins, both during and following seam narrowing (Fig. 5A,B). NMY-2 puncta also marked the transient medial AFB at the narrowest seam stage but did not appear to mark medial AFBs at later stages (Fig. 5A). Neither NMY-1 nor NMY-2 detectably marked hyp7 CFBs at the stages examined (n = at least 12 each). These data suggest that, at the time of seam narrowing, the longitudinal AFBs are part of a contractile actomyosin network. However, during seam relaxation, the medial AFBs (visible with UTRNCH::GFP) may no longer be contractile.

To test if NM II-dependent actomyosin constriction drives the observed changes in seam syncytium shape, we examined this tissue in NM II knockdowns at L4.4 (Fig. 5). Both *nmy-1(RNAi)* and *nmy-2(ts)* single mutants displayed misshapen syncytia with ectopic branches and protrusions (Fig. 5B). *nmy-1(RNAi)* larvae had seam widths only slightly larger than those of age-matched controls, while *nmy-2(ts)* single mutants (which show genetic interactions with the actin sensor - Fig. 2D) or *nmy-2(ts); nmy-1(RNAi)* double mutants had significantly distended seam syncytia, on average more than twice as wide as controls (Fig. 5B,C). Therefore, seam narrowing does depend on NM II.

To further characterize the relationship between NM II and the various actin structures observed, we examined the distribution of UTRNCH::GFP in NM II knockdowns. Junctional AFBs were still detected in *nmy-1(RNAi) nmy-2(ts)* double mutants but medial AFBs were largely fragmented or absent (Fig. 5C). These findings suggest that NM II is required for proper assembly of medial AFBs, implicating these structures in seam narrowing and alae shaping.

### Provisional matrix components are required for adult alae shaping but not seam narrowing

The misshapen or fragmented adult alae seen after actin, NM II, or junctional cadherin depletion are similar to those previously reported in some mutants affecting the provisional matrix that precedes each cuticle (Forman-Rubinsky et al., 2017; Gill et al., 2016). To test if other known components of this matrix are also required to pattern the adult alae, we used RNAi knockdown, mutant escapers, or mosaic approaches to circumvent the early arrest phenotypes seen in null mutants. RNAi knockdown or loss of the ZP proteins NOAH-1 or FBN-1 resulted in alae deformations similar to those previously reported after loss of the ZP protein LET-653 (Forman-Rubinsky et al., 2017) (Fig. 6A,B). Mosaic animals losing the lipocalin LPR-3 in subsets of seam cells (see Methods) also had regions of misshapen or fragmented alae and, in some cases, regions where alae were entirely missing (Fig. 6A,B). Therefore, the provisional matrix appears broadly important for forming and shaping adult alae.

**Figure 6.**
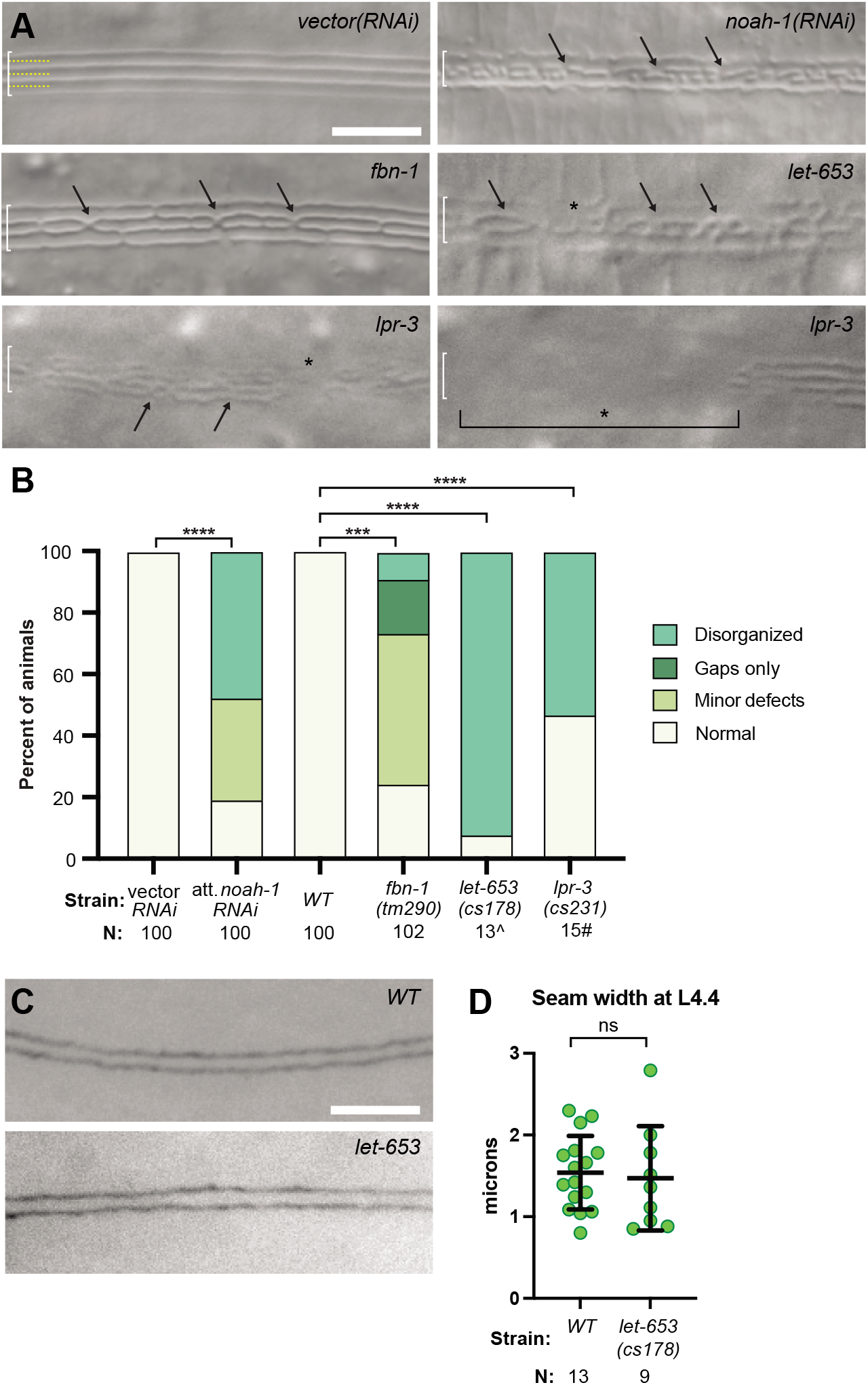
Provisional matrix components are required for patterning the alae. A) DIC images of alae in RNAi knockdowns or mutants of the indicated genotypes (strains N2, ARF251, 25°C; Strains UP3184, UP3452, 20°C). Arrows indicate disorganized regions. Asterisks indicate small breaks or gaps. Bracket indicates a large gap where no alae are present. B) Quantification of alae defects, as in Fig. 2D. ****P<0.0001, Fisher’s exact test. ^Data reproduced from (Forman-Rubinsky et al., 2017). #Numbers here are an under-estimate of the true penetrance of alae defects because not all mosaics would have lost *lpr-3* in the seam lineage (see Methods). H-J) At L4.4 stage, seam width (visualized with AJM-1::GFP) is similar between H) *WT* (strain SU93) and I) *let-653(cs178)* mutants (strain UP3184), both at 20°C. Measurements were performed as in Fig. 3B, and *WT* datapoints for the L4.4 stage are combined between both experiments. ns, P>0.05, Mann-Whitney test.

Sapio et al (2005) proposed that, in L1 and dauer larvae, ZP matrix polymerization might help generate the force that drives seam narrrowing. To test if ZP proteins in the provisional matrix are required for seam narrowing during the L4 stage, we examined *let-653* mutant L4s obtained by rescuing the lethal embryonic excretory tube defects with a tissue-specific rescue transgene (Methods). These mutants have defective alae (Fig. 6A,B) but are otherwise healthy. Seam morphology in these animals was examined with AJM-1::GFP. At L4.4, *let-653* mutants had fully fused and narrow seam cells that resembled those in wild-type (Fig. 6C,D), so at least this matrix component is not required for seam narrowing.

### Provisional matrices are patterned into longitudinal bands during and following seam narrowing

To examine the timing and patterns of provisional matrix deposition, we visualized the lipocalin LPR-3 and the ZP-domain proteins LET-653, FBN-1, and NOAH-1 using functional translational fusions expressed from the endogenous loci or from extrachromosomal transgenes (Cohen et al., 2020; Vuong-Brender et al., 2017) (Methods) (Fig. 7A). Provisional matrix proteins were visible over the seam syncytium during the L4.3-L4.4 stages, suggesting they are secreted prior to and/or during the period of seam narrowing (Fig. 7B Fig. 8A,C,E). The provisional matrix initially appeared diffuse rather than patterned, but over the next few hours, each protein resolved into a characteristic pattern of longitudinal bands that correspond to developing alae ridges or their associated valleys and borders (Fig. 7B-D). LPR-3 and LET-653 bands appeared quite early, before alae became clearly detectable by DIC. LPR-3 specifically marked three developing ridges, while LET-653 marked two or four valleys (Fig. 7B). NOAH-1 bands resolved slightly later and marked both ridges and valleys in different z-planes (Fig. 7B-D). FBN-1 was the last factor to become patterned, very close to the L4-Adult molt, and marked only valleys (Fig. 7B,D). Each protein also showed a different timeline of disappearance, but all disappeared from the alae region following the L4-adult molt. Therefore, these provisional matrix proteins are present during the period when alae are first being patterned and formed, but they are not permanent components of these ridge structures.

**Figure 7.**
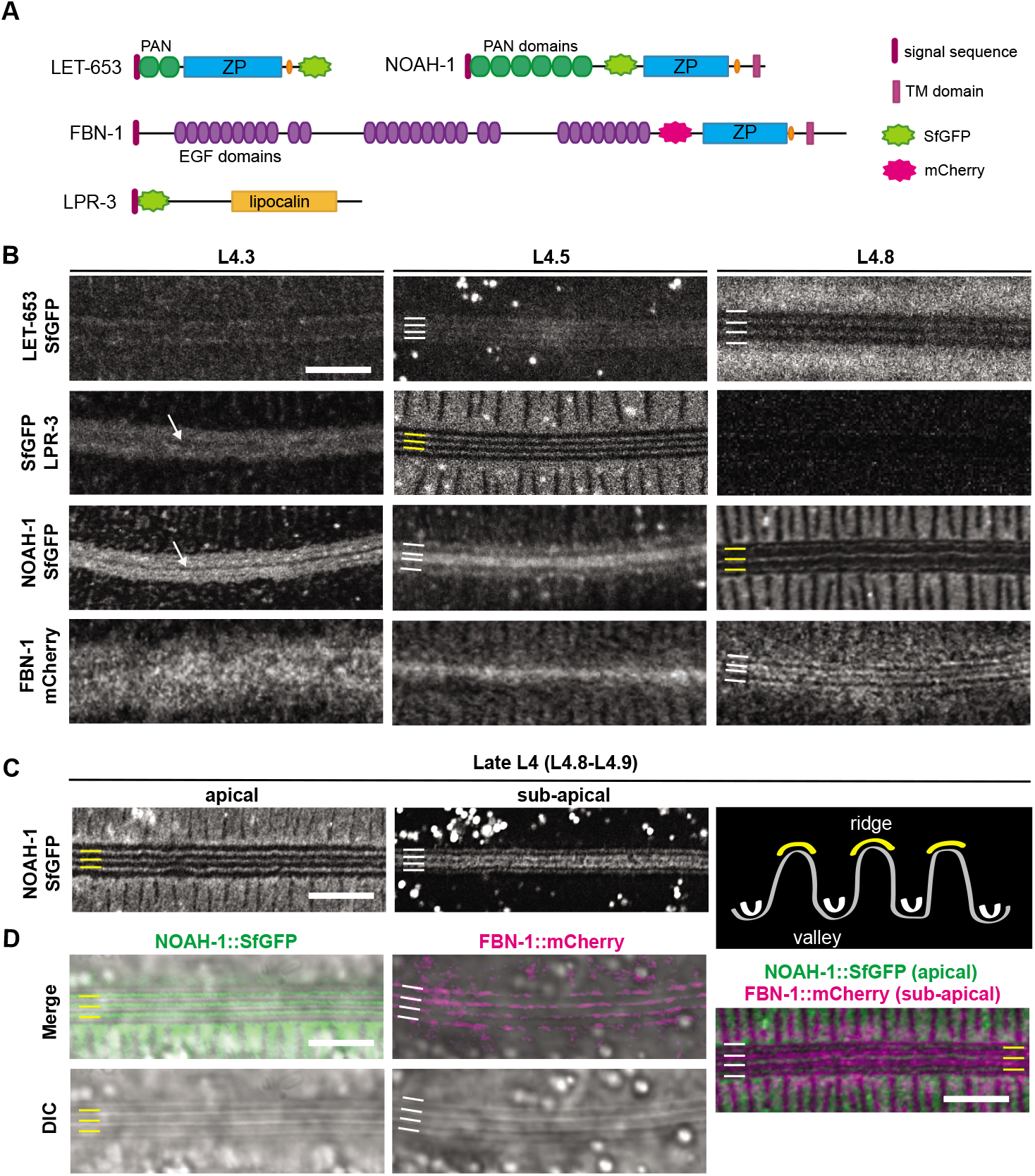
Provisional matrix patterns presage the ridges and valleys of adult-stage alae. A) Diagrams of provisional matrix components. B) Representative confocal slices show the dynamic distributions of indicated fusions in animals from mid-to late-L4. LET-653::SfGFP (strain UP3746, 20°C). SfGFP::LPR-3 (strains UP3666 or UP3693, 20°C). NOAH-1::SfGFP (strain ARF503 25°C). FBN-1::mCHERRY (strain ARF379, 25°C). C) Apical and sub-apical confocal slices from the same animal, showing the different NOAH-1::SfGFP patterns in different z-planes (strain CM10, 25°C). Some animals showed only one of these patterns, and others showed both simultaneously. Schematic shows interpretation of apical patterns as alae ridges and sub-apical patterns as valleys. Correspondingly, on all micrographs, yellow lines indicate developing alae ridges and white lines indicate flanking valleys. D) Airyscan-processed images of late L4s, showing that apical NOAH-1 aligns with alae ridges and sub-apical FBN-1 aligns with valleys, as seen by DIC. Image at right is a maximum-intensity projection showing FBN-1::mCHERRY and NOAH-1::SfGFP (strain ARF502 25°C) in different regions. All images are representative of at least n=5 per marker per stage. Scale bars: 5 μm.

**Figure 8.**
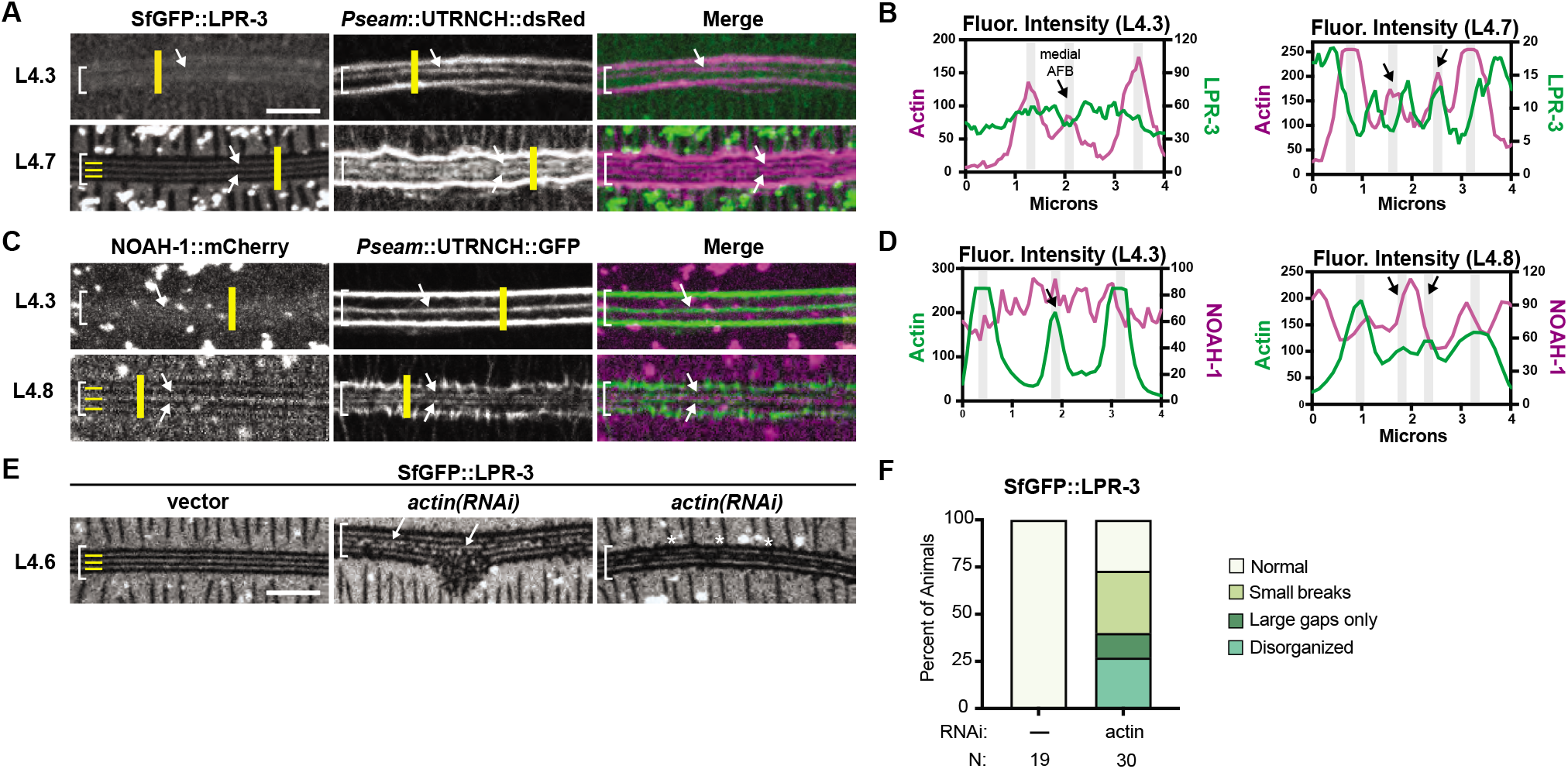
Actin is required to pattern the provisional matrix. A-D) Spatial relationships between seam AFBs and provisional matrix bands. A,C) Maximum intensity projections of animals expressing the designated actin sensor and matrix fusion. Brackets indicate seam region. Arrows point to medial AFBs. B,D) Line traces across the yellow bar region in raw versions of the corresponding images to left. Intensities for the two channels are plotted on separate scales as indicated. Grey shading represents the location of AFBs. Arrows point to medial AFB locations. A,B) SfGFP::LPR-3 apical bands are largely offset from AFBs (strain UP4127, 20°C). n=7 L4.3-L4.4. n= 8 L4.5-L4.7. The latter number includes only specimens where the UTRNCH::dsRed sensor detected medial AFBs in addition to junctional AFBs. C,D) NOAH-1::SfGFP apical bands are largely offset from AFBs (Strain UP4114, 20°C). n=6 L4.3-L4.4, n=10 L4.7-L4.8. E) *actin RNAi* disrupts provisional matrix patterns. Standard methods for bacterially-induced *actin RNAi* were used, and surviving animals were imaged at the L4.5-L4.7 stage (Strain UP3666, 20°C). F) Quantitation of SfGFP::LPR-3 patterns after actin depletion. Scale bars: 5 μm.

### Actin is required to pattern the provisional matrix into longitudinal bands

To understand the relationships between the seam longitudinal AFBs and the overlying longitudinal bands of provisional matrix, we combined our seam actin sensors with different matrix fusions (Fig. 8A-C). In narrowed seams, when three longitudinal AFBs were present, SfGFP::LPR-3 was cleared from a thin band directly overlying the medial AFB (Fig. 8A,B). A similar largely offset relationship was observed between the AFBs and the 3 apical bands of SfGFP::LPR-3 or NOAH-1::mCherry in later L4 animals (Fig. 8A-D). Together, these data suggest that AFBs underlie nascent valleys surrounding each developing ridge.

To test if AFBs might pattern the provisional matrix, we determined the effect of actin RNAi on LPR-3 matrix patterns. Following actin knockdown, SfGFP::LPR-3 often localized to labyrinthine deformities, rather than longitudinal bands (Fig. 8E,F). Furthermore, the remaining SfGFP::LPR-3 bands frequently contained many small breaks and regions that were faint and ill-defined (Fig. 8E,F). We conclude that cortical actin networks are required to pattern the provisional matrix into continuous longitudinal bands, and that loss or mis-patterning of the provisional matrix can explain mis-patterning of the permanent alae ridges.

### Ultrastructure of the epidermis reveals alternating sites of separation and adhesion among apical matrix layers

To characterize changes in the ultrastructure of the lateral epidermis and overlying apical matrices that happen while the alae take shape, we turned to transmission electron microscopy (TEM), using high pressure fixation to best preserve the fragile matrix (Hall et al., 2012; Weimer, 2006). We collected transverse sections through the mid-body of 10 distinct mid-L4 specimens, inspected the corresponding micrographs, and ordered the specimens by inference based on matrix appearance and comparisons to our DIC (Fig. 1D) and confocal (Figs. 3,4,7) image timelines. The micrographs shown in Figure 9 represent distinct steps in alae morphogenesis.

**Figure 9.**
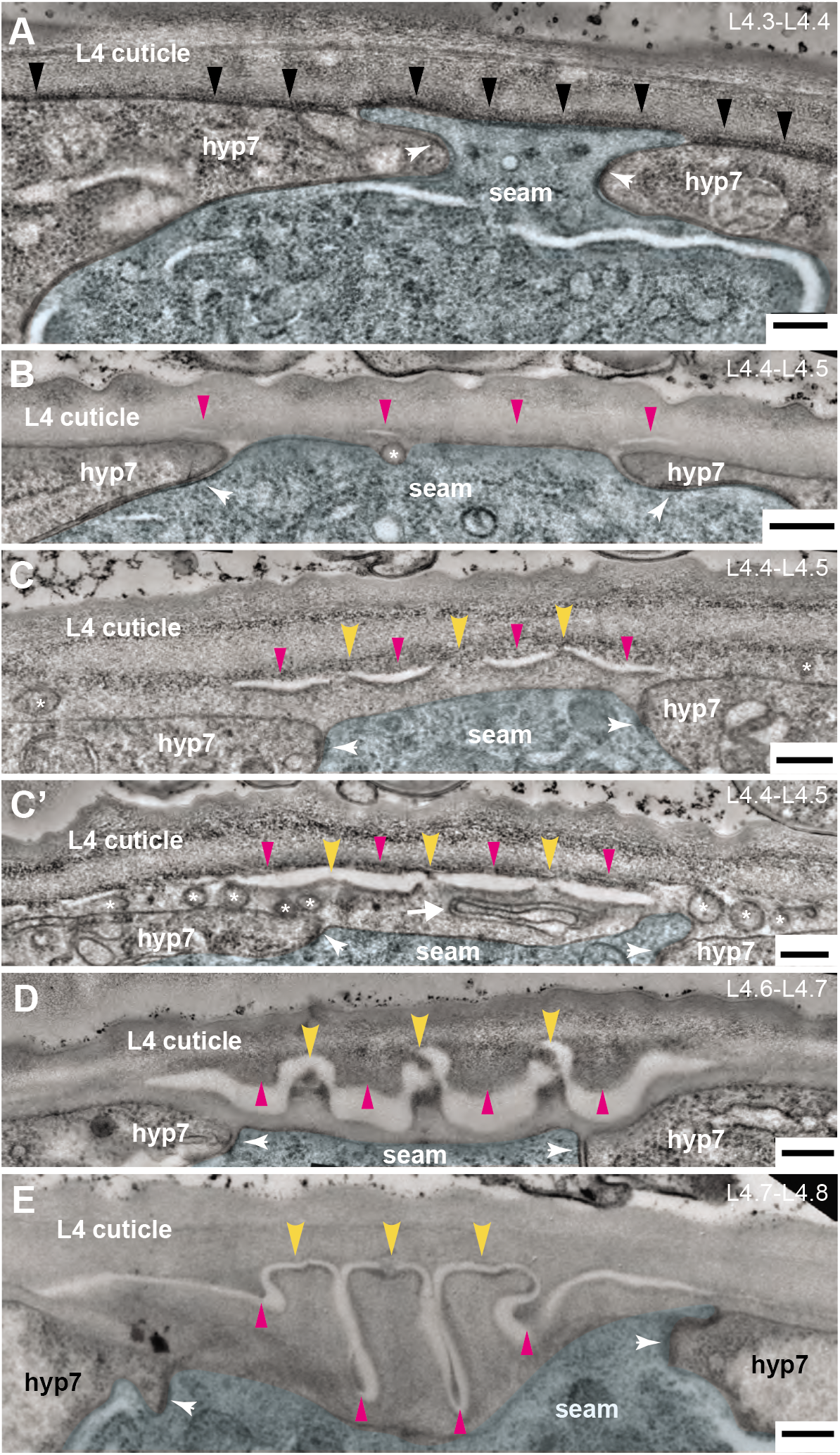
Ultrastructure of developing alae reveals differential matrix separation vs. adhesion. A-F) TEM micrographs of mid-to late-L4 *wild-type* (N2) specimens arranged by inferred age (total N=10). Transverse cuts through the mid-body are shown. See Fig. 1C for cartoon rendering of perspective. Seam cell is false colored in blue. White arrowheads indicate adherens junctions between the seam and hyp7 syncytia. Scale bars: 200 nm. A) ~L4.3-L4.4. The seam cell is highly constricted, with its narrowest point ~500nm in width. Black arrowheads indicate electron dense provisional matrix material on apical surfaces of both seam and hyp7 syncytia. B) ~L4.4-L4.5. Magenta arrowheads indicate four regions of provisional matrix separation. Asterisk marks a vesicle in transit across seam membrane. C) ~L4.4-L4.5. Yellow arrowheads indicate three regions of provisional matrix adhesion at nascent alae tips. Extracellular vesicles (asterisks) are present in the matrix over hyp7. C’) Regions of matrix separation are enlarged compared to panel C, which is another body region from the same specimen. Many extracellular vesicles (asterisks) and a larger membrane-bound structure (arrow) are present within the future adult cuticle. D) ~L4.6-L4.7. Discernable alae ridges have formed and contain electron dense material at their tips. Matrix fibrils connect these ridges to the L4 cuticle, while additional L4-cuticle-attached matrix protrudes down into the intervening gaps. E) ~L4.8. Maturing alae have grown in length and width, and valleys have narrowed. The central ridge still maintains a discernable connection to the L4 cuticle.

In two inferred L4.3-L4.4 stage specimens (Fig. 9A), the subapical region of the seam cell (at the adherens junctions) appears very narrow and pinched (0.5-1 micron wide), with hyp7 pushing in on both sides, consistent with apical constriction. Dark electron-dense extracellular material sits beneath the L4 cuticle, lining the apical plasma membrane of both the seam cell and hyp7; this material likely corresponds to the provisional matrix, which is deposited at this stage (Figs. 7,8). The seam cell apical membrane and L4 cuticle remain flat in these specimens. However, in a third specimen that also appears to be ~L4.4, the seam cell apical surfaces are still flat, yet the L4 cuticle has seemingly buckled into three deep folds (Fig. S1). This latter animal has a large break in the L4 cuticle at the vulva lumen, which may have released mechanical constraints on the tissue and matrix. Alternatively, cuticle buckling could be a normal but transient response to initial seam constriction.

In three specimens inferred to be just slightly older, about L4.4 to L4.5 stage (Fig. 9B-C’), the seam apical surface is narrow (0.9-1.1 microns) and the apical membrane and L4 cuticle are flat, but four discrete regions of separation appear between matrix layers over the seam. These separations define three intervening regions of remaining matrix adhesion, which we infer correspond to the future alae ridges. When comparing two images from different body regions in one of these specimens (Fig. 9C-C’), the separations are larger, and points of adhesion narrower, as ridges become more apparent. Dark electron-dense material lines the top and bottom surfaces of the separations, suggesting that both separations and adhesions occur between layers composed of the freshly deposited provisional matrix.

Another feature of these three specimens is the presence of many membrane-bound vesicles or organelles near the seam and hyp7 apical membranes, including in the apparent extracellular space where new adult cuticle is forming (Fig. 9B-C’). The extracellular vesicles (EVs) range in size from ~15 nm (the typical size of exosomes (Doyle and Wang, 2019)) to >600 μm (resembling migrasomes (Ma et al., 2015)). These EVs may contain materials for building or modifying the alae and cuticle.

In the four oldest specimens, inferred to range from L4.6 to L4.8, the seam cell is wider (1.5-3 microns) and contains many lamellar structures resembling lysosomes or lysosome-related organelles (LROs) (Fig. 9D,E; Fig. 10C). The nascent adult cuticle underneath the L4 cuticle shows progressively larger alae ridges, with deeper and narrower valleys separating the ridges. Small points of connection remain between these adult alae ridges and the thinning L4 cuticle above. Much additional matrix material has accreted at the base of the L4 cuticle and protrudes downward in a pattern that is complementary to that of the adult alae. This material is presumably “valley-localized” provisional matrix (Fig. 7C,D) that will be removed along with the old L4 cuticle at the molt.

**Figure 10.**
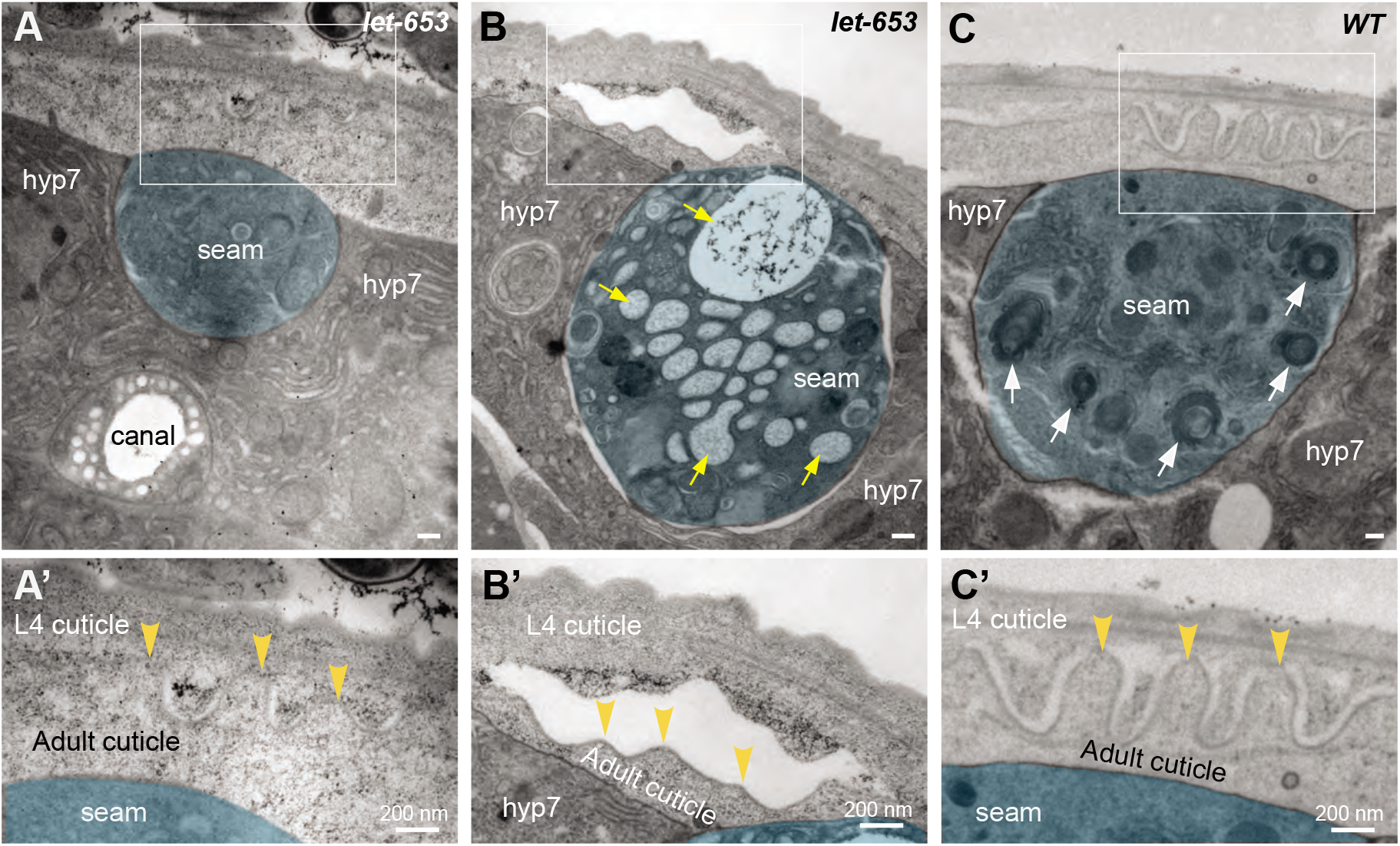
The provisional matrix component LET-653 is required for patterned adhesion versus separation of matrix layers. A-C) TEM micrographs of *let-653(cs178)* (A,B, strain UP3342) or wild-type (C, N2) L4 specimens. Transverse cuts through the mid-body are shown. Seam cell is false colored in blue. Scale bars: 200 nm. Boxed regions are shown at higher magnification in A’-C’. A, A’) There is no clear distinction of adhesive *vs*. separated matrix regions in this mid-L4 *let-653* specimen, despite the appearance of nascent alae ridges. B, B’) Mis-shapen alae ridges have completely separated from the L4 cuticle of this late L4 *let-653* specimen. Large vesicle structures (yellow arrows) fill the seam cell. C, C’) Many lysosomes or related lamellar organelles (white arrows) appear within the seam cell in wild-type late L4 specimens (N=4).

From these TEM data, we draw several key conclusions. First, many changes in matrix appearance occur immediately following the apex of seam narrowing. These changes are accompanied by the presence of vesicle populations, including EVs, that suggest active secretion by both the seam and hyp7 syncytia. Second, the seam apical membrane appears smooth throughout all stages of alae formation, with no evidence of upward protrusions or folds. Third, formation of alae involves differential separation vs. adhesion of matrix layers within the provisional matrix zone, with four regions of separation and three intervening regions of adhesion echoing the spacing of longitudinal AFBs within the relaxing seam and the banding patterns observed for provisional matrix proteins.

### Ultrastructure of the seam aECM in *let-653* mutants reveals requirements for the provisional matrix in patterning differential matrix adhesion

The above TEM data suggested that differential matrix separation occurs between layers of the provisional matrix. To better understand the contribution of provisional matrix to alae patterning, we also examined *let-653* mutant ultrastructure by TEM.

Figure 10 shows transverse TEM sections through the lateral epidermis of two mid-L4 *let-653* larvae processed identically and at the same time as the wild-type specimens above. In the first specimen, which appears to be the youngest of the two, developing alae ridges are discernible above the seam cell, but there are no clear distinctions between regions of matrix separation and adhesion (Fig. 10A). In the second specimen, the nascent alae cuticle has separated entirely from the above L4 cuticle (Fig. 10B). These data show that *let-653*, and likely the entire provisional matrix, is required for proper patterning of matrix separation vs. adhesion during alae formation. Thus, while some ridges do form in the mutants, they are not properly shaped or continuous.

The seam cell in the older *let-653* specimen also shows signs of abnormal protein trafficking, based on an accumulation of many large vesicles filled with amorphous fibrils characteristic of matricellular proteins (Fig. 10B). We previously noted and reported an accumulation of unusually large vesicles in vulF vulva cells of this same specimen (Cohen et al., 2020). In contrast, similar vesicles were not observed in seam syncytia of ten wild-type specimens, including four other late L4s (Fig. 10C). We conclude that, in addition to affecting matrix ultrastructure, the loss of *let-653* function interferes with the export or clearance of other matrix factors.

## DISCUSSION

Actomyosin networks and ZP-protein aECMs both play many roles in shaping epithelial tissues (Martin and Goldstein, 2014; Munjal and Lecuit, 2014; Plaza et al., 2010), but how the two systems are coordinated and how the actin cytoskeleton impacts complex aECM shapes remain poorly understood. Here we showed that actomyosin networks are required to pattern a provisional ZP-protein-rich matrix that determines the final structure of acellular alae ridges in the adult cuticle of *C. elegans*. Figure 11 summarizes our proposed model for how cytoskeletal patterns are translated into aECM patterns. The four key aspects of this model are: 1) Transient narrowing of the seam syncytium along the dorsal-ventral axis, through an atypical apical constriction mechanism involving longitudinal AFBs; 2) Transmission of AFB-dependent pulling forces across the seam apical membrane to a newly deposited, ZP-rich, provisional matrix, thereby generating local regions of separation between matrix layers; 3) Concurrent AFB-dependent patterning of the provisional matrix into longitudinal bands that pre-configure the final alae; 4) Recruitment of distinct collagens or other cuticle components to provisional matrix bands, thereby propagating regional differences in the short-lived provisional matrix to permanent differences in cuticle matrix structure. Below we discuss each of these aspects in more detail.

**Figure 11.**
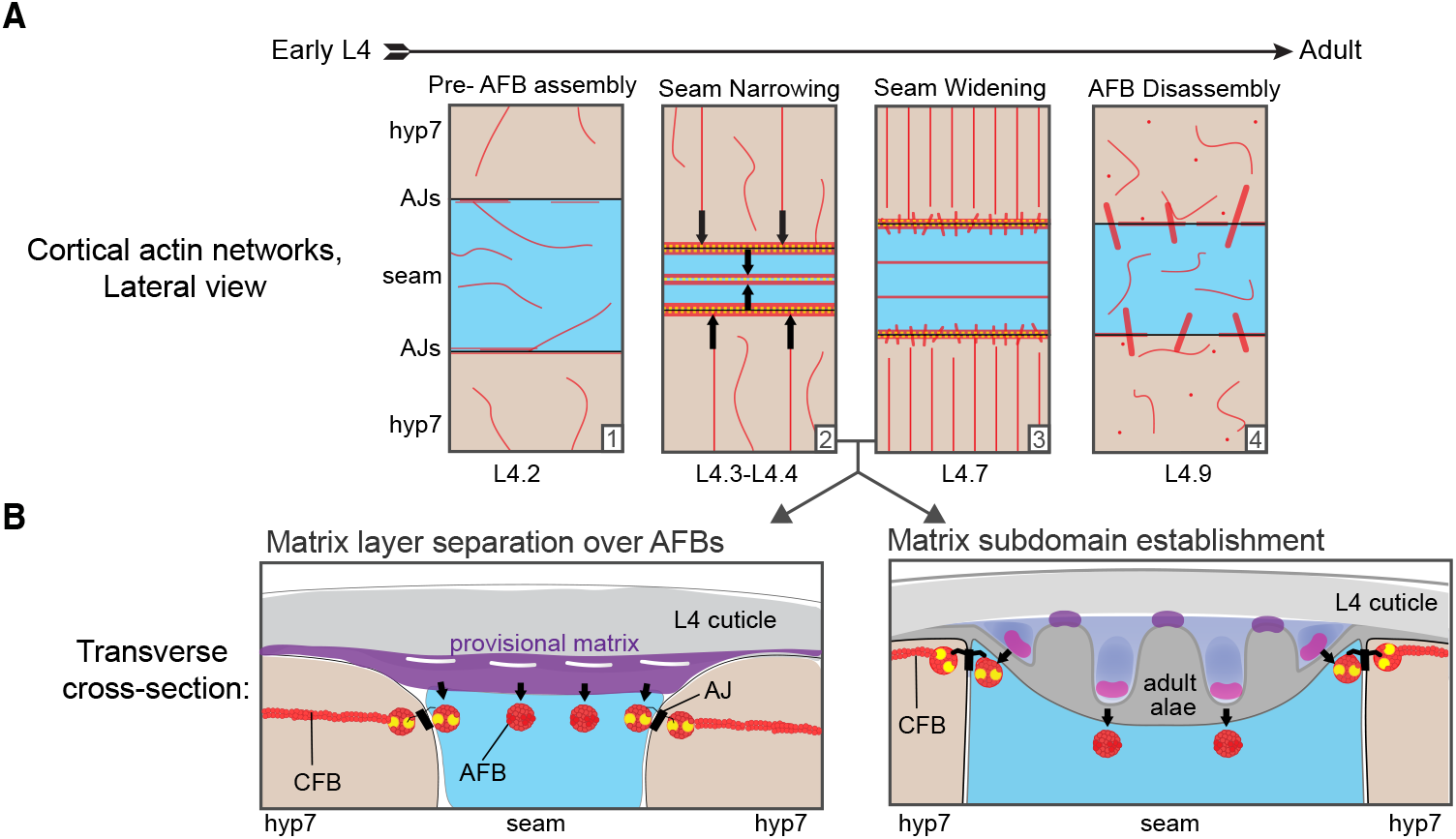
A mechanical relay model for development of the adult alae. Seam is blue, hyp7 peach. Cortical actin networks depicted in red, and NM II in yellow. **A)** Graphical representation of cortical actin networks across the larval-to-adult transition. Arrows indicate constrictive forces within the seam or pushing forces from hyp7 that lead to transient seam narrowing. (1) Epidermis prior to cortical actin network assembly; (2) Seam narrowing; (3) Seam relaxation. (4) Cortical actin network disassembly. **B)** Graphical representation of transverse cross-sections through the seam, hyp7, and overlying matrices during early (left) and late (right) stages of alae formation. (Early) Unknown connections link AFBs (red) to the overlying provisional matrix (purple). Pulling forces (arrows) from these connections generate small splits between molecularly distinct provisional matrix layers. (Late) As alae take shape, distinct provisional matrix components (purples, magenta) and cuticle components (grey) accumulate within different subdomains.

### An atypical apical constriction mechanism drives transient narrowing of the seam syncytium

Seam narrowing depends on NM II and occurs when actin and NM II localize to longitudinal AFBs flanking the AJs between the seam and hyp7. When attached to AJs, actomyosin filaments can generate pulling forces to constrict the apical surface of an epithelial cell (Martin and Goldstein, 2014; Munjal and Lecuit, 2014). Gastrulation, neurulation, and other essential morphogenetic processes depend on the apical constriction of particular cells within epithelial sheets (Goldstein and Nance, 2020; Martin and Goldstein, 2014; Martin et al., 2010). However, in these examples, AFBs typically are oriented isotropically or along the axis of tissue shortening (Koenderink and Paluch, 2018) rather than in parallel to it as we found in the seam. Instead, the organization of AFBs in the seam resembles that of AFBs within microridges, which are maze-like cellular protrusions on the epidermal surface of mucosal epithelia (Pinto et al., 2019; van Loon et al., 2020). In that case, NM II-containing minifilaments appear to connect parallel AFBs to generate constrictive forces that pull microridges closer together (van Loon et al., 2021). A similar mechanism could be operating during seam narrowing.

Seam narrowing also coincides with increased anisotropy of CFBs in hyp7 that appear to connect to the longitudinal AFBs and AJs. Local polymerization and upward extension of actin filaments can generate pushing forces at AJs (Efimova and Svitkina, 2018; Li et al., 2020; Papalazarou and Machesky, 2021). In theory, hyp7 CFBs could generate lateral pushing forces that also contribute to seam narrowing.

We found no evidence that forces associated with seam narrowing buckle the apical membrane to pattern ridge-like structures. In all ten TEM specimens examined, including those with very small, nascent alae, the seam plasma membrane appears smooth.

### Differential separation vs. adhesion of provisional matrix layers initiates alae formation

The provisional matrix is a temporary enclosure composed of several ZP-domain proteins (LET-653, NOAH-1, FBN-1), putative lipid transporters (LPR-3), and other still unknown matrix components that infiltrate beneath the cuticle. Many components of the provisional matrix are deposited on the apical surface of both the seam and hyp7 syncytia prior to and during the period of seam narrowing (Fig. 11B). Importantly, none of the matrix factors studied here are unique to the seam, yet they organize into alae-like structures only in this location.

Newly deposited matrix over the seam initially appears unpatterned, and actin is required for its subsequent organization into longitudinal bands (Fig. 11B). Ultrastructurally, the first sign of band patterning is the appearance of four evenly spaced, small separations between provisional matrix layers. The position of these initial matrix separations closely matches those of the four underlying seam AFBs present during seam relaxation. These separations will eventually define the valleys surrounding the adult alae, while the three intervening areas of remaining matrix adhesion define the adult alae ridges themselves. In the absence of LET-653, alae ridges *vs*. valleys appear poorly demarcated, consistent with a key role for the provisional matrix in subregion establishment.

### Relay of the seam AFB pattern to the provisional matrix

If the longitudinal AFBs do not push up on or buckle the seam apical membrane, then how are AFB patterns transmitted to the aECM? We propose that AFBs provide a downward pulling force that leads to separations in the matrix layers directly above them (Fig. 11B). This model requires that AFBs and the provisional matrix be somehow connected across the apical plasma membrane. In hyp7 at later stages, the actin cytoskeleton connects to cuticle aECM via matrilin-related proteins (Bercher et al., 2001; Hong et al., 2001). Distinct but analogous linkers may connect the actin cytoskeleton to the provisional aECM over the seam. This mechanical connection model does not need to invoke actomyosin contractility of the AFBs to explain the pulling mechanism; rather, as long as a physical connection is present, the motion of the worm’s body could generate sufficient tugging forces on the matrix overlying the AFBs.

A second, not mutually exclusive, possibility is that AFBs direct secretion of matrix proteases or other factors that modify the matrix at specific overlying sites. In Drosophila, NM II-mediated actomyosin contraction facilitates docking and compaction of large vesicles that deliver several aECM components to cell surfaces (Rousso et al., 2016), and cortical actin filaments are thought to specify sites of chitin synthase accumulation for chitin extrusion (Öztürk-Çolak et al., 2016). Longitudinal AFBs could serve a similar secretory role in the seam. However, we have not yet observed any evidence for patterned deposition of seam matrix factors or for patterned release of the EVs seen by TEM.

A key feature of the above models is that initially unpatterned matrix factors organize into longitudinal bands in response to an initiating cue from seam AFBs. Once the process is set in motion, self-organizing properties of the matrix may re-enforce the initial differences. For example, we propose that intrinsic biophysical properties could cause each protein to segregate preferentially toward or away from regions of tension, or to adhere preferentially to partners in different (now separated) layers, thereby establishing alternating matrix subregions with different molecular contents (Fig. 11B).

### The provisional matrix as a scaffold for permanent matrix assembly

Once the provisional matrix pattern is established, the next challenge is to grow and shape the alae ridges, while gradually replacing the provisional matrix with more permanent cuticle components such as collagens. *C. elegans* expresses more than 170 predicted cuticle collagen genes (Page and Johnstone, 2007), but so far very little is known about the collagen (or other matrix) content of adult alae. The cuticulins important for L1 or dauer alae formation do not seem to play a role in adults (Sapio et al., 2005). We hypothesize that binding interactions between specific provisional matrix factors and specific permanent matrix factors ultimately recruit different sets of factors to the alae ridges vs. valleys, thereby propagating the provisional matrix patterns to the permanent matrix.

Our TEM data suggest that alae enlargement is preceded by a burst of exocytosis that fills the developing matrix region over the seam with EVs of varying size. Seam EVs were previously suggested to release hedgehog-related cargos (Liegeois et al., 2006), and we hypothesize that EVs could also be a major route by which other types of matrix cargo or proteases are delivered to the alae. Many of the EVs appear to derive from hyp7, so one role of seam narrowing could be to facilitate dumping of contents from hyp7 into a narrow extracellular space over the seam. Some of the larger EVs resemble migrasomes (Ma et al., 2015) and might simply be debris left behind by hyp7 as it begins to withdraw during seam relaxation. Once more permanent alae components are identified, it will be interesting to test if they are trafficked though EVs and to investigate how the processes of matrix delivery, infiltration, and replacement occur.

*A* key conclusion from this study is that provisional matrix proteins can play critical roles in shaping a permanent aECM structure even when these proteins are not present in that final structure. Provisional matrices likely are important intermediates in forming and shaping other permanent aECM structures too. For example, in Drosophila, a temporary luminal matrix precedes formation of a distinct chitin-rich cuticle within tracheal tubes (Ozturk-Colak et al., 2015). A dramatic example of a provisional matrix in mammals is that of tooth enamel, which initially contains many matrix proteins, most of which are removed and replaced by calcium phosphate during the process of mineralization (Moradian-Oldak and George, 2021). In cases where the final aECM structure must be rigid enough to retain its shape, it makes sense to pattern a more malleable provisional matrix first and then to use that matrix as a scaffold or mold on which to build the permanent structure.

## MATERIALS AND METHODS

### Strains and Transgenes

*C. elegans* strains used in this study are listed in Table S1. Strains were maintained under standard conditions (Brenner, 1974) and cultivated at either 20°C or 25°C, as indicated in Figure legends. For synchronization, gravid worms were allowed to lay eggs during limited time windows or were bleached to isolate eggs and hatchlings arrested in starvation-induced L1 diapause and were released from diapause by plating on NGM seeded with *E. coli* OP50-1. The precise stages of L4 worms were further determined based on the shape of the vulva (Cohen et al., 2020; Mok et al., 2015).

Strains with conditional *nmy-2* alleles were propagated at permissive temperature (15°C) and cultivated at restrictive temperature (25°C) following release from starvation-induced L1 diapause. *let-653* mutant defects were examined in animals rescued to viability with a transgene driven by the excretory duct-specific promoter *lin-48* (Johnson et al., 2001). *lpr-3* seam mosaics were obtained from transgenic strain UP3452 *[lpr-3(cs231); csEx436 [lpr-3+; myo-2::mCherry]*; potential mosaics were recognized based on loss of the transgene-associated *myo-2::mCherry* signal in one or more of the three pharyngeal muscle 3 (pm3) cells, two of which derive from the ABa lineage and thus are lineally-related to seam cells (Sulston and Horvitz, 1977). 10/15 such mosaics had alae defects, compared to only 1/21 non-mosaic siblings (p=0.0001, Fisher’s Exact Test).

Table S2 describes the oligonucleotides used in this study. Phusion High Fidelity Polymerase (NEB) was used to amplify DNA for sequencing and cloning. Gibson assembly (NEB) and standard cloning reactions were used to construct fusion genes and corresponding plasmids. To create the *egl-18p::rde-1*+ *egl-18p::rde-1::sl2::mCherry::unc-54* 3’UTR fusion gene housed in pSK08, the promoter of *egl-18*, which corresponds to nucleotides 1910072-1913471 of chromosome IV (GenBank: NC_003282); the coding region of *rde-1*, which corresponds to nucleotides 9988043-9991614 of chromosome V (GenBank: NC_003283); coding sequence for *mCherry* (GenBank: KT175701), *sl2* (GenBank: LK928133); and the *unc-54* 3’UTR cassette from pPD95.75 were combined. To construct the *dpy-7p::rde-1*+ *dpy-7p::rde-1::sl2::nls-gfp::unc-54* 3’-UTR fusion gene housed in pSK38, the minimal promoter of *dpy-7*, which corresponds to nucleotides 7537789-7537869 and 7537914-7538219 of chromosome X (GenBank: NC_003284); the coding region of *rde-1; SL2;* and the *nls-gfp::unc-54* 3’UTR cassette from pPD95.73 were united. To construct the *dpy-7p::utrnch::dsRed::unc-54* 3’UTR fusion gene housed in pSK26, the promoter of *dpy-7* (as above); the coding sequence for the first CH domain (residues 1-261) of human Utrophin (GenBank: LX69086); the coding sequence for *dsRed* (GenBank: HQ418395); the *unc-54* 3’ UTR cassette from pPD95.81; and the pUC57 backbone were combined. To construct the *egl-18p::utrnch::gfp::unc-54* 3’ UTR fusion gene housed in pSK34, the promoter of *egl-18;* the sequence encoding UTRNCH; and the *gfp::unc-54* 3’ UTR cassette and backbone from pPD95.81 were united. All variants of plasmid pPD95 were gifts from Andy Fire.

To construct the *noah-1::sfGFP::noah-1* translational fusion gene housed in pCM05, regulatory and coding regions of *noah-1* were amplified from genomic DNA (nucleotides 5874389-5883950 of chromosome I, GenBank: LK927608) and cloned into a NotI-filled derivate of pCR-Blunt II-TOPO (Invitrogen). A *NotI-sfgfp-NotI* cassette was inserted in-frame between the codons for P624 and V625 of *noah-1a* (Genbank: NM_170870). The corresponding NotI site was created using a Q5 mutagenesis kit (Invitrogen). Superfolder (sf) GFP was isolated from pCW11 (Max Heiman, Harvard University).

All extrachromosomal arrays were generated by microinjection of young adults with mixtures containing 100ng/μl DNA. To generate *aaaEx37*, pSK08 (5ng/μl), *ttx-3::gfp* (40ng/μl), and pRS316 (55ng/μl) were co-injected into JK537 *rde-1(ne219)*. To generate *aaaEx162*, pSK38 (5ng/μl), *ttx-3::dsred* (40ng/μl) and pRS316 were co-injected into JK537. To generate aaaEx108, pSK26 (0.5ng/μl), *ttx-3::gfp*, and pRS316 were co-injected into N2. To generate *aaaEx117*, pSK34 (5ng/μl), *ttx-3::gfp*, and pRS316 were co-injected into N2. Optimal plasmid concentrations used to generate tandem utrnch arrays were empirically determined by titration. UTRNCH signals were readily detected in the resulting transgenic animals, while phenotypes associated with high levels of UTRN were not observed. To generate *aaaEx78 [fl-fbn-1::mCherry::fbn-1]*, pSK27 (2.5 ng/μl), the above-mentioned PCR product (1.15 ng/μl), *ttx-3p::gfp*, and pRS316 were co-injected into N2. To generate *aaaEx167*, pCM05 (1ng/μl), *ttx-3::dsred*, and pRS316 were co injected into ARF379. Resulting transgenic lines were out-crossed to N2 to remove *aaaIs12*. The extrachromosomal arrays *aaaEx78* and *aaaEx167* rescued lethality caused by respective null alleles of *fbn-1* or *noah-1*, confirming the production of functional fusion proteins. Extrachromosomal arrays were integrated into the genome by UV irradiation at 450 kJ using an FB-UVXL-1000 (Fisher Scientific). Strains with newly integrated arrays were back crossed to JK537 or N2 4 to 6 times prior to further analyses.

### RNA-mediated interference (RNAi)

Bacterial-mediated RNAi was performed as described (Kamath et al., 2003), except that NGM (nematode growth medium) plates were supplemented with 8mM rather than 1mM, isopropyl β-D-1-thiogalactopyranoside (IPTG). For attenuated RNAi treatments, animals were washed off from control plates 14hrs after release from L1 diapause with 14 ml M9, rotated for 30 minutes in M9 to remove residual gut bacteria and then transferred to experimental RNAi plates. As a control, worms were fed the same *E. coli* HT115(DE3) transformed with the empty vector pPD129.36. Upon induction by IPTG, such bacteria produce short dsRNA molecules that do not match any annotated gene of *C. elegans*.

To knock down *actin* by bacterial-mediated RNAi, we used a sequence-verified clone for *act-2* present in the Ahringer library (Kamath et al., 2003). To knock down *nmy-1* (Genbank: LK927643), 1121 bp of genomic DNA from exon 10 was cloned into pPD129.36, the standard expression vector for dsRNAs. For *zoo-1* (GenBank: NM_001026515), cDNA spanning exons 1–7 was cloned into pPD129.36, as previously described (Lockwood et al., 2008). For *noah-1* (GenBank: LK927608), 1024bp from exon 6 was cloned into pPD129.36. Each of the resulting plasmids (pSK43, pSK44 and pCM13) was verified by Sanger sequencing and used to transform *E. coli* strain HT115(DE3).

### DiI Staining of Cuticles

DiI staining to visualize cuticle structures was performed essentially as described (Schultz and Gumienny, 2012). Briefly, approximately 600 adult worms were incubated in 400 μl of 30 μg/mL DiI (Sigma) in M9 for 3 hours, shaking at 350 rpm. Worms were then washed 1X in M9 buffer, re-suspended in 100 μl of M9, and dispensed to a 6-cm NGM plate seeded with *E. coli* OP50-1. To remove excess unbound dye, worms were allowed to crawl on the plate for 30 minutes prior to imaging.

### Microscopy and Image Analyses

Worms were anesthetized with sodium azide (2.5%) and/or levamisole (10 mM) in M9 buffer and mounted on 2% agarose pads. A Zeiss Axioplan or Axioskop microscope (Carl Zeiss Microscopy) with an attached Hamamatsu Orca ER CCD camera or Leica DFC360 FX camera was used for compound microscopy. Images were acquired and analyzed using the software packages Volocity 6.3 (PerkinElmer) or Qcapture (Qimaging). The confocal images in Figs. 2, 4, 5C, were captured on a Zeiss LSM5 controlled by ZEN 9.0 software. Images in Fig. 3 and the NOAH-1 and FBN-1 images in Fig. 7, were captured on a Zeiss LSM880 with Airyscan processing; the dual colors image in Fig. 7D and Fig. 8C were captured on the same instrument with spectral analysis and signals from dsRed and GFP then linear unmixed in Zen 9.0. Confocal images of LET-653 and LPR-3 in Fig. 7B were captured with Leica TCS SP8 confocal microscope. Confocal images in Figs. 5A,B and 8A,C were captured with a Leica DMi8 confocal microscope. Image intensity and color were adjusted in FIJI or Photoshop for presentation.

Alae defects were quantified using a scoring rubric that prioritized regions of disorganized alae over gaps where alae were completely missing; animals that displayed each phenotype in different body regions were placed in the former category.

Measurements were made using Volocity 6.3 (PerkinElmer), ImageJ (Version 1.48v, NIH), and Fiji (Schindelin et al., 2012). Seam width was measured in Fiji as the distance between AJM-1-marked junctions; six measurements at 100-point intervals were made per image and averaged. For NM II knockdown experiments, where seam shape was too abnormal for standard width measurements, normalized seam width was calculated based on seam area (surface area within AJM-1::mCHERRY boundaries, measured in Volocity), divided by the distance imaged along the A-P axis. The ImageJ plugin FibrilTool was used to measure CFB anisotropy {Boudaoud, 2014 #3}. For each worm assayed, 6 values were obtained by subdividing the lateral region of hyp7 into 3 dorsal and 3 ventral ROIs, each approximately 400 μm^2^. Line scans in Fig. 8 were performed on raw images using a 10pt line and the PlotProfile tool in FIJI.

For transmission electron microscopy (TEM), synchronized wild-type (N2) and *let-653* mutant (UP3342) L4 animals were collected and processed by high pressure freezing followed by freeze substitution into osmium tetroxide in acetone (Hall et al., 2012; Weimer, 2006). Specimens were rinsed and embedded into LX112 resin and cut transversely to generate thin sections of approximately 70 nm each. At least two sections each from two different mid-body regions were sampled per animal. Sections were stained with uranyl acetate and lead citrate and observed on a JEM-1010 (Jeol, Peabody Massachusetts) transmission electron microscope. Images were processed in ImageJ and manually pseudocolored in Adobe Illustrator (Adobe, San Jose California). We imaged a total of n=10 N2 and n=2 UP3342 specimens.

### Statistical Analyses

GraphPad Prism 6 and Microsoft Excel 15.21 were used for statistical analyses. In all dot-plots or bar graphs, lines and error bars indicate the mean and standard deviation, and dots represent mean measurements from individual animals. To perform statistical analyses on categorical data, phenotypic categories were combined such that outcomes were classified as abnormal versus superficially normal (Figs. 2 and 6).

## ACKNOWLEDGEMENTS

Some strains used in this study were provided by the *Caenorhabditis* Genetics Center, which is funded by the NIH Office of Research Infrastructure Programs (P40 OD010440). We are grateful to Margot Quinlan, Alvaro Sagasti, Larry Zipursky, Ron Ellis, and members of our laboratories for helpful discussions. We also thank Helen Schmidt and Nick Serra for critical reading of the manuscript, and Michel Labouesse, Margot Quinlan, John Kim, Eric Miska, and David Sherwood for sharing reagents. This work was supported by the National Science Foundation (IOS1258218 to ARF), the National Institutes of Health (R01GM125959 and R35GM136315 to MVS, OD010943 to DHH, and training awards GM007185 to SK and T32 AR007465 to JDC), and by a Dissertation Year Fellowship from the UCLA Graduate Division (to SK).

## SUPPLEMENTAL MATERIALS

**Supplemental Figure 1. Outlier wild-type L4 TEM specimen with broken and buckled cuticle**

Cuticle breaks may release mechanical tension and allow buckling of matrix over the seam. Seam is false colored in blue. Boxes indicate regions shown in panels to right. A, A’) TEM section through a mid-body region far from the vulva. The cuticle over the seam is arranged into deep folds. We interpret this cuticle to be that of the mid-L4 stage. Note mature appearance of the cuticle, small seam cell size, and absence of receding cuticle or lamellar LROs typically associated with late L4s (compare to sibling specimens in Figs. 9 and 10). B, B’) TEM section through the vulva region of the same specimen. A large break in the L4 cuticle is present at the vulva, which has a large lumen characteristic of mid-L4s and no signs yet of adult cuticle formation. Note connection of dorsal vulva cells to the seam and distortion of seam cell shape in this region. Scale bars: 5 μm.

**Supplemental Table 1:** *C. elegans* strains used in this study.

**Supplemental Table 2:** Oligonucleotides used in this study.

## References

Baird, G. S., Zacharias, D. A. and Tsien, R. Y. (2000). Biochemistry, mutagenesis, and oligomerization of DsRed, a red fluorescent protein from coral. Proc. Natl. Acad. Sci. U. S. A. 97, 11984–11989.

Bercher, M., Wahl, J., Vogel, B. E., Lu, C., Hedgecock, E. M., Hall, D. H. and Plenefisch, J. D. (2001). mua-3, a gene required for mechanical tissue integrity in Caenorhabditis elegans, encodes a novel transmembrane protein of epithelial attachment complexes. J. Cell Biol. 154, 415–426.

Brenner, S. (1974). The genetics of Caenorhabditis elegans. Genetics 77, 71–94.

Burkel, B. M., von Dassow, G. and Bement, W. M. (2007). Versatile fluorescent probes for actin filaments based on the actin-binding domain of utrophin. Cell Motil. Cytoskeleton 64, 822–832.

Cohen, J. D. and Sundaram, M. V. (2020). C. elegans Apical Extracellular Matrices Shape Epithelia. J. Dev. Biol. 8, E23.

Cohen, J. D., Flatt, K. M., Schroeder, N. E. and Sundaram, M. V. (2019). Epithelial Shaping by Diverse Apical Extracellular Matrices Requires the Nidogen Domain Protein DEX-1 in Caenorhabditis elegans. Genetics 211, 185–200.

Cohen, J. D., Sparacio, A. P., Belfi, A. C., Forman-Rubinsky, R., Hall, D. H., Maul-Newby, H., Frand, A. R. and Sundaram, M. V. (2020). A multi-layered and dynamic apical extracellular matrix shapes the vulva lumen in Caenorhabditis elegans. eLife 9, e57874.

Costa, M., Draper, B. W. and Priess, J. R. (1997). The role of actin filaments in patterning the Caenorhabditis elegans cuticle. Dev. Biol. 184, 373–84.

Costa, M., Raich, W., Agbunag, C., Leung, B., Hardin, J. and Priess, J. R. (1998). A putative catenin-cadherin system mediates morphogenesis of the Caenorhabditis elegans embryo. J Cell Biol 141, 297–308.

Cox, G. N. (1981). The cuticle of Caenorhabditis elegans II: Stage-specific changes in ultrastructure and protein composition during postembryonic development. Dev Biol 86, 456–470.

Dickinson, D. J., Ward, J. D., Reiner, D. J. and Goldstein, B. (2013). Engineering the Caenorhabditis elegans genome using Cas9-triggered homologous recombination. Nat. Methods 10, 1028–34.

Ding, S. S. and Woollard, A. (2017). Non-muscle myosin II is required for correct fate specification in the Caenorhabditis elegans seam cell divisions. Sci. Rep. 7, 3524.

Doyle, L. M. and Wang, M. Z. (2019). Overview of Extracellular Vesicles, Their Origin, Composition, Purpose, and Methods for Exosome Isolation and Analysis. Cells 8, E727.

Efimova, N. and Svitkina, T. M. (2018). Branched actin networks push against each other at adherens junctions to maintain cell-cell adhesion. J. Cell Biol. 217, 1827–1845.

Fernandes, I., Chanut-Delalande, H., Ferrer, P., Latapie, Y., Waltzer, L., Affolter, M., Payre, F. and Plaza, S. (2010). Zona pellucida domain proteins remodel the apical compartment for localized cell shape changes. Dev Cell 18, 64–76.

Flatt, K. M., Beshers, C., Unal, C., Cohen, J. D., Sundaram, M. V. and Schroeder, N. E. (2019). Epidermal Remodeling in Caenorhabditis elegans Dauers Requires the Nidogen Domain Protein DEX-1. Genetics 211, 169–183.

Forman-Rubinsky, R., Cohen, J. D. and Sundaram, M. V. (2017). Lipocalins Are Required for Apical Extracellular Matrix Organization and Remodeling in Caenorhabditis elegans. Genetics 207, 625–642.

Gaudette, S., Hughes, D. and Boller, M. (2020). The endothelial glycocalyx: Structure and function in health and critical illness. J. Vet. Emerg. Crit. Care San Antonio Tex 2001 30, 117–134.

Gill, H. K., Cohen, J. D., Ayala-Figueroa, J., Forman-Rubinsky, R., Poggioli, C., Bickard, K., Parry, J. M., Pu, P., Hall, D. H. and Sundaram, M. V. (2016). Integrity of Narrow Epithelial Tubes in the C. elegans Excretory System Requires a Transient Luminal Matrix. PLoS Genet. 12, e1006205.

Gilleard, J. S., Barry, J. D. and Johnstone, I. L. (1997). cis regulatory requirements for hypodermal cell-specific expression of the Caenorhabditis elegans cuticle collagen gene dpy-7. Mol. Cell. Biol. 17, 2301–11.

Goldstein, B. and Nance, J. (2020). Caenorhabditis elegans Gastrulation: A Model for Understanding How Cells Polarize, Change Shape, and Journey Toward the Center of an Embryo. Genetics 214, 265–277.

Hall, D. H., Hartwieg, E. and Nguyen, K. C. Q. (2012). Modern electron microscopy methods for C. elegans. Methods Cell Biol. 107, 93–149.

Hannezo, E., Dong, B., Recho, P., Joanny, J.-F. and Hayashi, S. (2015). Cortical instability drives periodic supracellular actin pattern formation in epithelial tubes. Proc. Natl. Acad. Sci. U. S. A. 112, 8620–8625.

Harland, D. P. and Plowman, J. E. (2018). Development of Hair Fibres. In The Hair Fibre: Proteins, Structure and Development (ed. Plowman, J. E.), Harland, D. P.), and Deb-Choudhury, S.), pp. 109–154. Singapore: Springer.

Hong, L., Elbl, T., Ward, J., Franzini-Armstrong, C., Rybicka, K. K., Gatewood, B. K., Baillie, D. L. and Bucher, E. A. (2001). MUP-4 is a novel transmembrane protein with functions in epithelial cell adhesion in Caenorhabditis elegans. J Cell Biol 154, 403–14.

Johansson, M. E., Sjovall, H. and Hansson, G. C. (2013). The gastrointestinal mucus system in health and disease. Nat Rev Gastroenterol Hepatol 10, 352–61.

Johnson, A. D., Fitzsimmons, D., Hagman, J. and Chamberlin, H. M. (2001). EGL-38 Pax regulates the ovo-related gene lin-48 during Caenorhabditis elegans organ development. Development 128, 2857–65.

Kamath, R. S., Fraser, A. G., Dong, Y., Poulin, G., Durbin, R., Gotta, M., Kanapin, A., Le Bot, N., Moreno, S., Sohrmann, M., et al. (2003). Systematic functional analysis of the *Caenorhabditis elegans* genome using RNAi. Nature 421, 231–237.

Kang, H., Bang, I., Jin, K. S., Lee, B., Lee, J., Shao, X., Heier, J. A., Kwiatkowski, A. V., Nelson, W. J., Hardin, J., et al. (2017). Structural and functional characterization of Caenorhabditis elegans α-catenin reveals constitutive binding to β-catenin and F-actin. J. Biol. Chem. 292, 7077–7086.

Katsanos, D., Ferrando-Marco, M., Razzaq, I., Aughey, G., Southall, T. D. and Barkoulas, M. (2021). Gene expression profiling of epidermal cell types in C. elegans using Targeted DamID. Dev. Camb. Engl. 148, dev199452.

Knight, C. G., Patel, M. N., Azevedo, R. B. R. and Leroi, A. M. (2002). A novel mode of ecdysozoan growth in Caenorhabditis elegans. Evol. Dev. 4, 16–27.

Koenderink, G. H. and Paluch, E. K. (2018). Architecture shapes contractility in actomyosin networks. Curr. Opin. Cell Biol. 50, 79–85.

Koh, K. and Rothman, J. H. (2001). EGL-18 and ELT-6 are required continuously to regulate epidermal seam cell differentiation and cell fusion in C. elegans. Dev. Camb. Engl. 128, 2867–2880.

Koppen, M., Simske, J. S., Sims, P. A., Firestein, B. L., Hall, D. H., Radice, A. D., Rongo, C. and Hardin, J. D. (2001). Cooperative regulation of AJM-1 controls junctional integrity in *Caenorhabditis elegans* epithelia. Nat. Cell Biol 3, 983–991.

Lehrbach, N. J., Armisen, J., Lightfoot, H. L., Murfitt, K. J., Bugaut, A., Balasubramanian, S. and Miska, E. A. (2009). LIN-28 and the poly(U) polymerase PUP-2 regulate let-7 microRNA processing in Caenorhabditis elegans. Nat. Struct. Mol. Biol. 16, 1016–1020.

Li, J. X. H., Tang, V. W. and Brieher, W. M. (2020). Actin protrusions push at apical junctions to maintain E-cadherin adhesion. Proc. Natl. Acad. Sci. U. S. A. 117, 432–438.

Liegeois, S., Benedetto, A., Garnier, J. M., Schwab, Y. and Labouesse, M. (2006). The V0-ATPase mediates apical secretion of exosomes containing Hedgehog-related proteins in Caenorhabditis elegans. J. Cell Biol. 173, 949–61.

Lloyd, V. J. and Nadeau, N. J. (2021). The evolution of structural colour in butterflies. Curr. Opin. Genet. Dev. 69, 28–34.

Lockwood, C., Zaidel-Bar, R. and Hardin, J. (2008). The C. elegans zonula occludens ortholog cooperates with the cadherin complex to recruit actin during morphogenesis. Curr Biol 18, 1333–13337.

Ma, L., Li, Y., Peng, J., Wu, D., Zhao, X., Cui, Y., Chen, L., Yan, X., Du, Y. and Yu, L. (2015). Discovery of the migrasome, an organelle mediating release of cytoplasmic contents during cell migration. Cell Res. 25, 24–38.

Martin, A. C. and Goldstein, B. (2014). Apical constriction: themes and variations on a cellular mechanism driving morphogenesis. Dev. Camb. Engl. 141, 1987–1998.

Martin, A. C., Gelbart, M., Fernandez-Gonzalez, R., Kaschube, M. and Wieschaus, E. F. (2010). Integration of contractile forces during tissue invagination. J. Cell Biol. 188, 735–749.

Melak, M., Plessner, M. and Grosse, R. (2017). Actin visualization at a glance. J. Cell Sci. 130, 525–530.

Mok, D. Z., Sternberg, P. W. and Inoue, T. (2015). Morphologically defined sub-stages of C. elegans vulval development in the fourth larval stage. BMC Dev Biol 15, 26.

Moores, C. A. and Kendrick-Jones, J. (2000). Biochemical characterisation of the actin-binding properties of utrophin. Cell Motil. Cytoskeleton 46, 116–128.

Moradian-Oldak, J. and George, A. (2021). Biomineralization of Enamel and Dentin Mediated by Matrix Proteins. J. Dent. Res. 100, 1020–1029.

Munjal, A. and Lecuit, T. (2014). Actomyosin networks and tissue morphogenesis. Dev. Camb. Engl. 141, 1789–1793.

Muriel, J. M., Brannan, M., Taylor, K., Johnstone, I. L., Lithgow, G. J. and Tuckwell, D. (2003). M142.2 (cut-6), a novel Caenorhabditis elegans matrix gene important for dauer body shape. Dev. Biol. 260, 339–351.

Ozturk-Colak, A., Moussian, B. and Araujo, S. J. (2015). Drosophila chitinous aECM and its cellular interactions during tracheal development. Dev Dyn.

Öztürk-Çolak, A., Moussian, B., Araújo, S. J. and Casanova, J. (2016). A feedback mechanism converts individual cell features into a supracellular ECM structure in Drosophila trachea. eLife 5, e09373.

Page, A. P. and Johnstone, I. L. (2007). The cuticle. In Wormbook (ed. The C. elegans Research Community),.

Papalazarou, V. and Machesky, L. M. (2021). The cell pushes back: The Arp2/3 complex is a key orchestrator of cellular responses to environmental forces. Curr. Opin. Cell Biol. 68, 37–44.

Pasti, G. and Labouesse, M. (2014). Epithelial junctions, cytoskeleton, and polarity. Wormbook 1–35.

Piekny, A. J., Johnson, J. L., Cham, G. D. and Mains, P. E. (2003). The Caenorhabditis elegans nonmuscle myosin genes nmy-1 and nmy-2 function as redundant components of the let-502/Rho-binding kinase and mel-11/myosin phosphatase pathway during embryonic morphogenesis. Development 130, 5695–704.

Pinto, C. S., Khandekar, A., Bhavna, R., Kiesel, P., Pigino, G. and Sonawane, M. (2019). Microridges are apical epithelial projections formed of F-actin networks that organize the glycan layer. Sci. Rep. 9, 12191.

Plaza, S., Chanut-Delalande, H., Fernandes, I., Wassarman, P. M. and Payre, F. (2010). From A to Z: apical structures and zona pellucida-domain proteins. Trends Cell Biol 20, 524–32.

Podbilewicz, B. and White, J. G. (1994). Cell fusions in the developing epithelial of C. elegans. Dev. Biol. 161, 408–24.

Priess, J. R. and Hirsh, D. I. (1986). Caenorhabditis elegans morphogenesis: the role of the cytoskeleton in elongation of the embryo. Dev Biol 117, 156–73.

Qadota, H., Inoue, M., Hikita, T., Koppen, M., Hardin, J. D., Amano, M., Moerman, D. G. and Kaibuchi, K. (2007). Establishment of a tissue-specific RNAi system in C. elegans. Gene 400, 166–73.

Rousso, T., Schejter, E. D. and Shilo, B.-Z. (2016). Orchestrated content release from Drosophila glue-protein vesicles by a contractile actomyosin network. Nat. Cell Biol. 18, 181–190.

Sapio, M. R., Hilliard, M. A., Cermola, M., Favre, R. and Bazzicalupo, P. (2005). The Zona Pellucida domain containing proteins, CUT-1, CUT-3 and CUT-5, play essential roles in the development of the larval alae in Caenorhabditis elegans. Dev Biol 282, 231–45.

Schaeffer, C., Devuyst, O. and Rampoldi, L. (2021). Uromodulin: Roles in Health and Disease. Annu. Rev. Physiol. 83, 477–501.

Schindelin, J., Arganda-Carreras, I., Frise, E., Kaynig, V., Longair, M., Pietzsch, T., Preibisch, S., Rueden, C., Saalfeld, S., Schmid, B., et al. (2012). Fiji: an open-source platform for biological-image analysis. Nat. Methods 9, 676–82.

Schultz, R. D. and Gumienny, T. L. (2012). Visualization of Caenorhabditis elegans cuticular structures using the lipophilic vital dye DiI. J Vis Exp 59, e3362.

Sebastiano, M., Lassandro, F. and Bazzicalupo, P. (1991). cut-1 a Caenorhabditis elegans gene coding for a dauer-specific noncollagenous component of the cuticle. Dev Biol 146, 519–30.

Sellon, J. B., Ghaffari, R. and Freeman, D. M. (2019). The Tectorial Membrane: Mechanical Properties and Functions. Cold Spring Harb. Perspect. Med. 9, a033514.

Singh, R. N. and Sulston, J. E. (1978). Some observations on molting in C. elegans. Nematologica 24, 63–71.

Sulston, J. E. and Horvitz, H. R. (1977). Post-embryonic cell lineages of the nematode *Caenorhabditis elegans*. Dev Biol 56, 110–156.

van Loon, A. P., Erofeev, I. S., Maryshev, I. V., Goryachev, A. B. and Sagasti, A. (2020). Cortical contraction drives the 3D patterning of epithelial cell surfaces. J. Cell Biol. 219, e201904144.

van Loon, A. P., Erofeev, I. S., Goryachev, A. B. and Sagasti, A. (2021). Stochastic contraction of myosin minifilaments drives evolution of microridge protrusion patterns in epithelial cells. Mol. Biol. Cell 32, 1501–1513.

Vuong-Brender, T. T. K., Suman, S. K. and Labouesse, M. (2017). The apical ECM preserves embryonic integrity and distributes mechanical stress during morphogenesis. Development.

Weimer, R. M. (2006). Preservation of C. elegans tissue via high-pressure freezing and freeze-substitution for ultrastructural analysis and immunocytochemistry. Methods Mol. Biol. 351, 203–21.

Whitsett, J. A., Wert, S. E. and Weaver, T. E. (2015). Diseases of pulmonary surfactant homeostasis. Annu Rev Pathol 10, 371–93.

Willis, J. H., Munro, E., Lyczak, R. and Bowerman, B. (2006). Conditional dominant mutations in the Caenorhabditis elegans gene act-2 identify cytoplasmic and muscle roles for a redundant actin isoform. Mol Biol Cell 17, 1051–64.

Zhang, H., Landmann, F., Zahreddine, H., Rodriguez, D., Koch, M. and Labouesse, M. (2011). A tension-induced mechanotransduction pathway promotes epithelial morphogenesis. Nature 471, 99–103.

