## Supplementary figures and images for "A transient apical extracellular matrix relays cytoskeletal patterns to shape permanent acellular ridges on the surface of adult *C. elegans*"

### Supplemental Figure 1

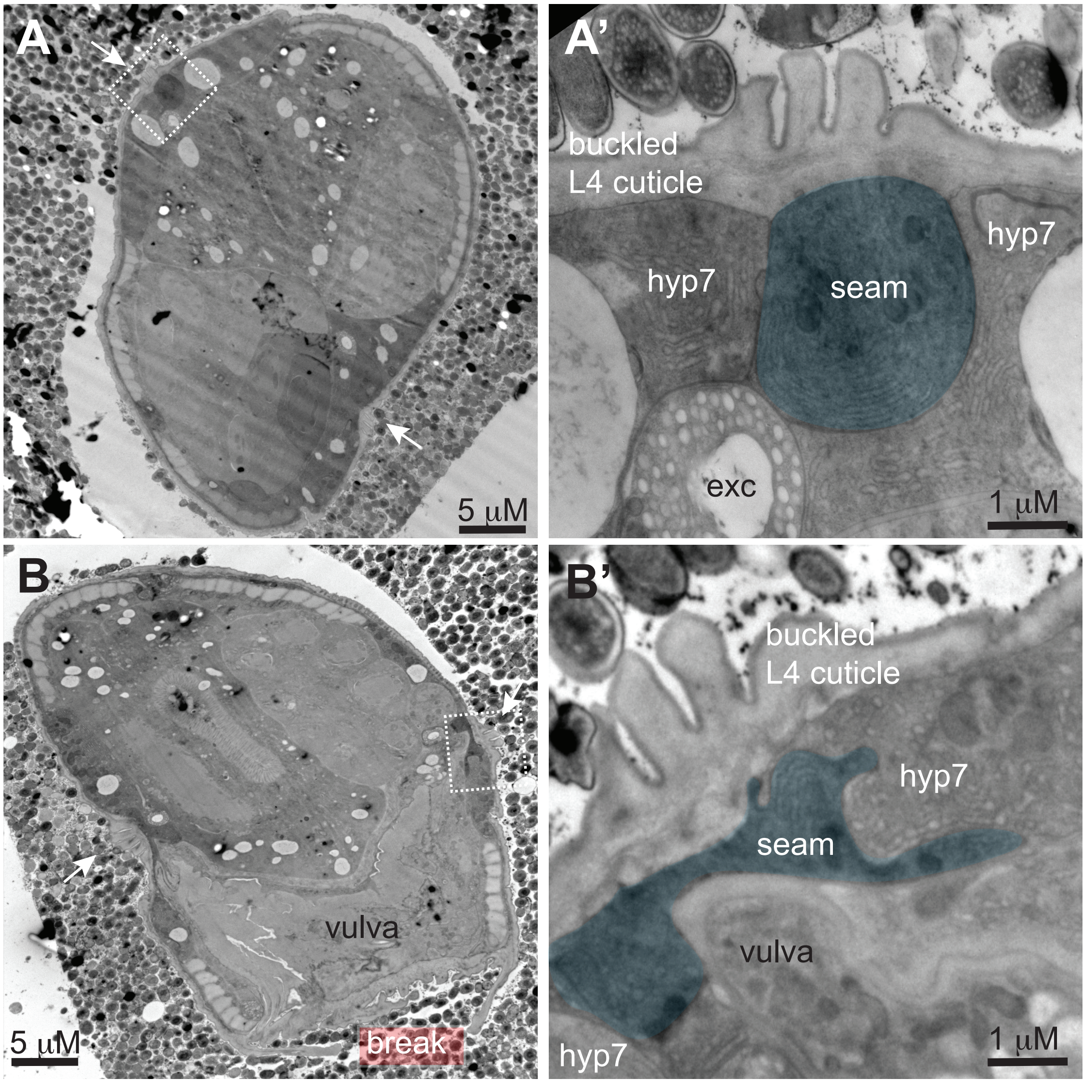
